# Pump-less, recirculating organ-on-a-chip (rOoC) platform

**DOI:** 10.1101/2022.09.06.506239

**Authors:** M. Busek, A. Aizenshtadt, T. Koch, A. Frank, L. Delon, M. Amirola Martinez, A. Golovin, C. Dumas, J. Stokowiec, S. Gruenzner, E. Melum, S. Krauss

**Author notes:** Authors contributed equally to this work (M. Busek, A. Aizenshtadt).

## Abstract

Current organ-on-a-chip (OoC) systems mimic important aspects of specific organ and tissue functions, however, many commercial and academic devices are either too simple for advanced assays or require a complicated support set-up including external driving systems such as pumps and tubing that hamper scalability and robustness.

We have developed a novel, pump-less directional flow recirculating organ-on-a-chip (rOoC) platform that creates continuous or pulsed directional gravity-driven flow by a combination of a 3D-tilting system and an optimized microfluidic layout.

The rOoC platform allows growing and connecting tissue or organ representations on-chip with the possibility of incorporating barrier functions, gradients, and circulating cells. Using the rOoC platform we demonstrate simple and reproducible endothelialisation, hepatic organoid integration, and the first steps of vascularization of 3D organ representations on-chip.

## Introduction

Organ-on-a-Chip (OoC) technology combines microfluidics and cell biology to mimic tissue or organ functionality *in vitro*. OoCs are on the verge of widespread use as new models with significant advantages for studying diseases and drug response/toxicity in both academia and the pharmaceutical industry^1^. A variety of platforms were developed from 2010 onwards when the first complex OoC, a Lung-on-a-Chip model comprising a co-cultivation of lung alveolar and endothelial cells (ECs) on the two sides of a porous membrane, was presented ^2^. A major driver for OoC development is the pharmaceutical industry^3^. Bringing a new drug to the market can cost more than 1 billion US$^4^ and OoC technology is expected to reduce R&D costs by 10–30%^5^. Another important aspect of OoC technology is *in vitro* disease modelling^6^ as human physiology frequently differs from laboratory animal physiology. In addition, pathological processes may be controlled and monitored on-chip. Finally, OoC technology is an important step toward reducing animal testing in line with the 3R initiative^7^.

However, after more than 10 years of development, OoC technology is still limited in its applicability and scalability. This might be caused by the necessity of most formats to use expensive and complicated driving/pumping systems thus limiting long-term stability and usability for smaller laboratories.

Recently, we surveyed peers in academia to understand the major limitations, needs and advantages of OoC technology^8^. We identified major problems in terms of usability, and robustness which is in line with surveys conducted by others^9^. Moreover, we found that many scientists aim for more complex *in vitro* models for their studies than those that are currently available on user-friendly platforms. Hence, important features of new OoCs generation include among others the incorporation of blood vessel-like structures, 3D culture models and immune cells. Existing OoCs tend to combine these sought-after features at the cost of low scalability and usability^10–13^.

Here we present a novel, scalable, and easy-to-use OoC platform, termed rOoC that not only creates a directed flow without the need for pumps and tubing but also allows the implementation of biological complexity on-chip addressing several of the concerns raised above. The rOoC platform supports on-chip vascularization, growth factor and nutrient gradients, 3D organoids culture and the incorporation of circulating cells. We present the layout and fabrication of the rOoC platform, its technical characterization, and a mathematical model to predict the flow in the device under different conditions. Next, we demonstrate stable EC culture over 2 weeks of directional flow-dependent endothelial alignment and sprouting of microvessels into the organoid compartment (OC) of the chip. We then show the long-term viability and functionality of 3D liver organoids on-chip. Finally, we demonstrate that the rOoC platform is a suitable platform also for circulating immune cells. The device can support several perfusion and OCs that are separated by an extracellular matrix (ECM) barrier, a layout that can in principle be applied to a plethora of single and combined organ models.

## Results

### 1) Platform design and fabrication

Adapted from common gravity-driven perfusion systems placed on a 2D-tilting platform^14^, we introduce a novel concept of using a 3D-tilting/rotating platform (Fig. 1a) to produce a directed recirculating flow on our rOoC platform without using pumps. To generate uni-directional flows, we propose the usage of elongated and high reservoirs (Fig. 1/2a). By connecting two reservoirs with microchannels, a fluidic ring circuit is produced and allows the cyclic filling and emptying of the reservoirs hence producing a directed flow as shown in the Suppl. Video 1. The main effect responsible for the directionality of the flow is that the liquid level in the reservoirs equalizes during tilting thus causing the inlet to run dry and thus preventing backflow. For actuation, we use a standard laboratory rotator (3D rotator waver, VWR Avantor, USA). The flow rate in the rOoC depends on the volume (both maximum and filled with liquid) and geometry of the reservoirs, the perfusion channel (PC) dimensions, as well as the tilt and rotation speed of the platform.

**Figure 1:**
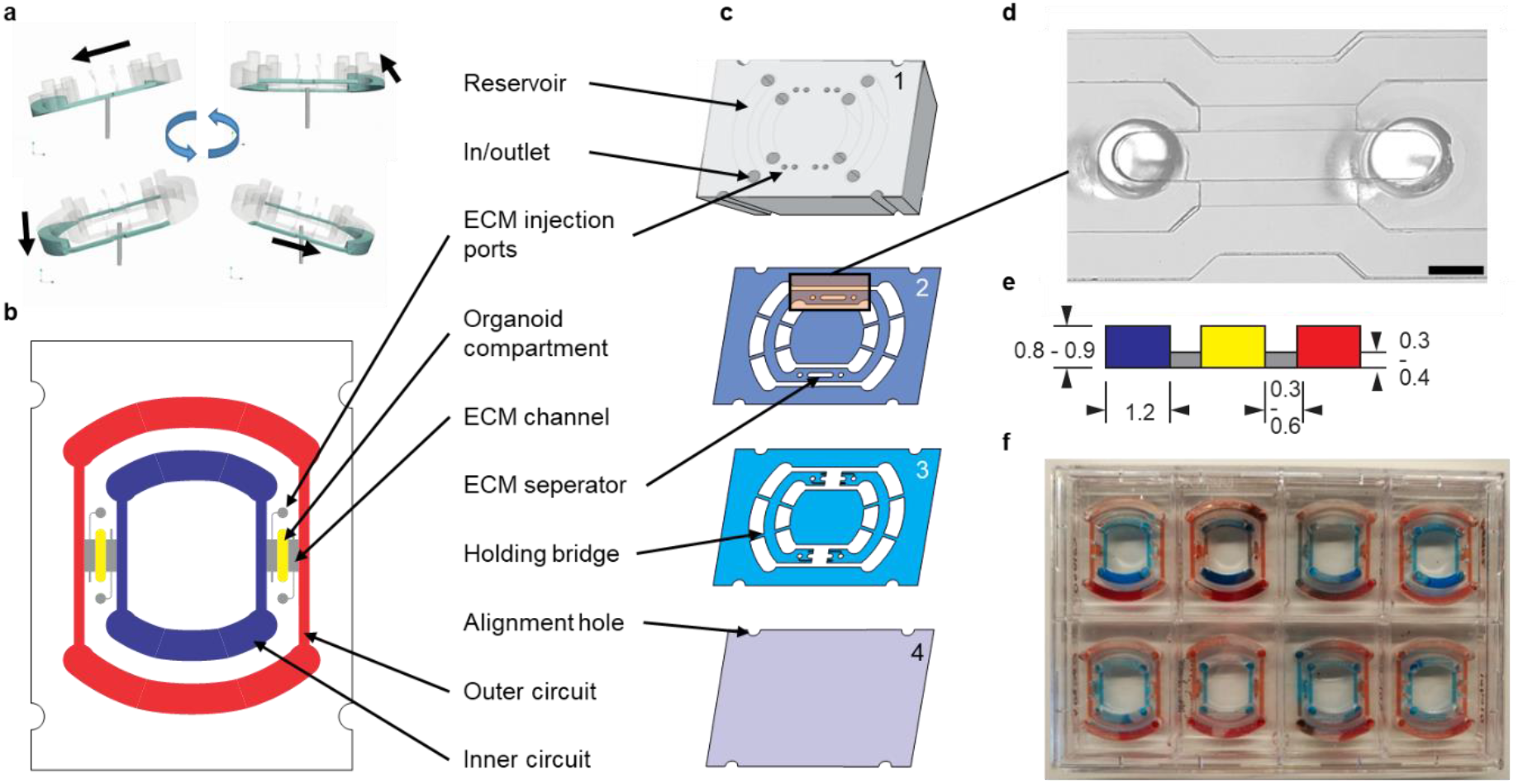
Platform layout. a) Fluid movement principle based on a 3D-tilting motion. b) Schematic view of the rOoC device indicating the two circuits (outer: red, inner: blue), the organoid compartments (OC) (yellow) and the ECM channels (grey). c) Explosion view. d) Microscopic image of the OC and the perfusion channels (PC). Scale bar: 1 mm. e) Cross-section of the stepped channel geometry with dimensions used in this study (dimensions in mm). f) 8 chips placed in a rectangular 8-well plate.

**Figure 2:**
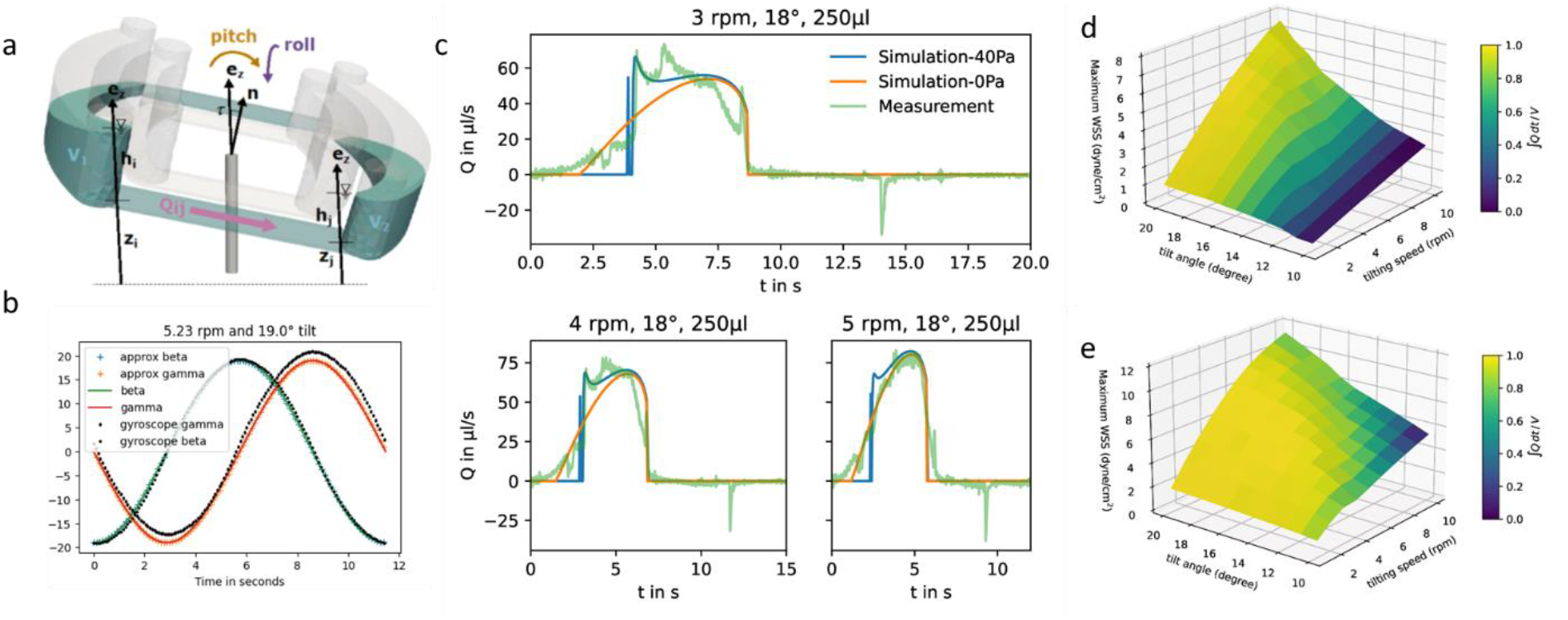
Flow characterization and modelling. a) Graphical explanation of used symbols. b) Measured and calculated pitch and roll (beta/gamma) of the platform. c) Modelled vs. measured flow velocities (Q) for different tilting speeds at 18° tilt and 250 μL volume (green - μPIV measurement, orange/blue – simulation with 0/40 Pa critical pressure). Maximum WSS for different tilts and speeds in the inner (d) and outer (e) PC for a filling volume of 200/300 μL. The colour of the plot indicates directionality. For ∫Q/V< 1 the flow is bi-directional.

The PC dimensions are based on the sizes of smaller arteries in the body^15^. We use a cross-section of 1.2 × 0.8 mm^2^. To have a media-to-cell ratio comparable to standard cell culture plastics (96/48 well plates), we selected a reservoir volume of 200 μL for the inner and 300 μL for the outer circuit. The reservoir shape was designed half-round as this geometry is less prone to trapping circulating cells. The general chip layout (Fig. 1b) follows the commonly used three-lane concept as this is especially useful for vascularization studies^16^. Our unique actuation principle gives us the possibility to place the two PCs within each other with two OCs positioned in between (Fig. 1b). We selected a length of 4 mm, a width between 0.5 and 1.2 mm and a height of 0.8/0.9 mm for the OCs as this is big enough to house several organoids (between 200-500 μm each) and to allow endothelial sprouting into the OC. For loading, we mixed cells or organoids with ECM and injected the mixture into the OC through the two ECM injection ports (diameter: 1mm, accessible by a 10 μL-pipette tip). When the ECM solidifies, the organoids are fixed in place.

The rOoC is built up of 4 layers of Poly(methyl methacrylate) (PMMA) (Fig. 1c). The layers are designed in a CAD program, laser-cut, precisely stacked and thermally bonded together^17,18^. The process is detailed in the methods part. The 4 layers are:

1. The top layer (6 mm) includes laser-engraved reservoirs and ECM injection ports.
2. Layer 2 is 0.5 mm thick and includes the reservoirs, PCs, OCs and connecting holes for the ECM-cell mixture.
3. Layer 3 includes PCs and OCs and is 0.4 mm thick. In contrast to layer 2, a connection between the PCs and OCs is included that can be filled with ECM.
4. The rOoC is sealed with a thin film of PMMA (0.175 mm).

Together, layers 2 and 3 form a stepped channel layout (Fig. 1d)that prevents ECM from entering the PCs^19^. Due to capillary forces, the liquid ECM is retained in the region where the channel height is abruptly increased. By using this layout, a biodegradable separation layer^20^ is formed between the OCs and the PCs allowing for spatial defining of the cell composition^19^ and studying cellular interaction such as angiogenesis^21^. The connection area is 0.3-0.6 mm wide so that ECs can sprout into the OC while ensuring that ECM is not accidentally blocking the PCs. Additional separate ECM channels can be used to fill the ECM (possibly mixed with other cells) independently from the OCs. For on-chip confocal imaging, only PMMA films with superior optical clarity were chosen. To improve scalability, up to 8 rOoCs can be placed in a standard rectangular 8-well plate (Thermo Scientific, catalogue No. 267062) and covered with a lid (Fig. 1e). In this configuration, the evaporation was measured (by weighing) to be 31 ± 16 μL per day for a total filling volume of 500 μL (n=6).

The configuration of the rOoC with two nested circuits of PCs separated by ECM channels and OCs allows filling the two circuits with different media to create a gradient in the OC which can be maintained in the ECM by dosing reagents in either the outer or the inner circuit. In many biological processes, including differentiation, vascularization and functional subdivisions such as liver zonation, growth factor-, nutrient- and gas gradients play a crucial role and need to be included when representing tissues or organs. To test the capability of the rOoC device to establish and maintain gradients, 70kDa fluorescein-labelled Dextran was filled in the inner PC and its gradient within the OC was observed under a fluorescence microscope. The detailed setup and experimental data are given in Appendix A. The dextran gradient was produced and assessed to be stable over 6 hours in the rOoC. Note that the gradient formation and characteristics will differ between molecules and used ECM due to different molecular weights and interactions between the ECM and the permeating substances (binding to ECM).

### 2) Flow characterisation and modelling

For ECs cultivated at the PC walls, the wall shear stress (WSS) is an important parameter for cell functionality. We have developed a mathematical model to predict flow rates and WSS values in the rOoC and cross-validated it with micro-particle velocimetry measurements (μPIV). From the flow velocity v, the WSS can be calculated from the gradient of the tangential velocity in the normal direction to the wall:

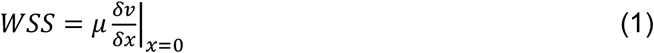

The 3D-tilting motion of the platform can mathematically be described as the rotation (frequency f) of a normal vector 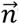that is tilted by an angle of α to the z-axis (Fig. 2a). This motion leads to a time-dependent pitch and roll (β, γ) of the platform. We measured both values with a gyroscope (Arduino 9-Axis Motion Shield coupled to an Arduino Uno, Arduino, Ivrea, Italy - source code see Suppl. Code 1). The data was recorded on a computer and saved as CSV-File. A plot of β and γ is given in Fig. 2b. Mathematically, β and γ can be described using trigonometric functions and the rotation function ϑ(*t*) = 2*πft* (a detailed explanation is given in Appendix B):

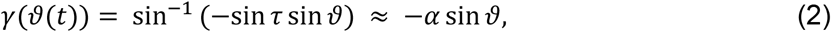

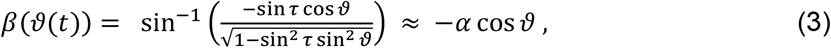

As shown in Fig. 2b, the approximation has an error of less than 1% for a tilt of 20° or less and thus can be used to describe γ and β. The Python script used to generate the plot is given in the Suppl. Code 2.

The computational model (detailed in the Methods part, source code in Suppl. Code 3) allows for calculating the flow rates dynamically for different reservoir geometries, tilt angle, angular velocities, and filling volumes. Changing channel geometry requires the re-computation of the channel transmissibility and relations between flow rate and velocity or WSS. The following parameters have been used for the computational model:

The calculation yields the flow-time curve given in Fig. 2c that aligns well with the measured flow-time curves (flow measurement described in the materials and methods part). We calculated the flow-time curve with two critical pressure values (0 and 40 Pa) to investigate the influence of the surface tension on the flow characteristics. It seems that for low tilting speeds (< 4 rpm) a better fitting to the measured data can be obtained for 40 Pa critical pressure whereas for speeds >5 rpm surface effects can be neglected and the curve aligns very well with the prediction using 0 Pa critical pressure.

A possible explanation for this effect is evaporation. Due to the high-intensity LED illumination that is required for the μPIV measurement, the evaporation is significantly higher than in the incubator and might cause the liquid to evaporate fully in the reservoirs when they run dry. This is especially true for lower tilting speeds as the remaining liquid in the emptied reservoir has more time to evaporate.

The calculated flow profile (see Suppl. Text) indicates that the flow velocity is maximum in the middle of the channel and reaches zero near the walls. In contrast, the WSS (plot see Suppl. Text) is maximum at the middle of each wall but zero at the corners. We next calculated the maximum WSS with the parameters given in Table 1 (viscosity μ = 0.8 mPa*s for T=37 °C) for different tilts and rotation speeds for both the outer and the inner PC (*V*_*res*_ = 200 *μL*, *l*_*c*_ = 12 *mm*) and plotted these values in Fig. 2d/e. The colour in the plots indicates whether the whole reservoir volume is transported thus indicating if the flow is uni-directional 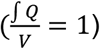. If this quotient becomes <1 a backflow occurs and if this quotient equals zero, the flow is fully bi-directional similar to 2D tilting systems (no net flow in one direction). In the rOoC, this is for instance the case for low tilts (<10°), high rotations speed (>10 rpm) and also predominantly in the inner PC. Moreover, bi-directional flow occurs for higher filling volumes (Appendix C).

**Table 1:** Parameters used for computational modelling and flow measurements

| Expression | $b_c$ | $w_c$ | $l_{c,out}$ | $V_{max}$ | $p_c$ | $\alpha$ | $f$ | $\mu$ | $P$ |
| --- | --- | --- | --- | --- | --- | --- | --- | --- | --- |
| Name | PC width | PC height | PC Length | Reservoir volume | Critical pressure | Tilt | Tilting speed | Fluid viscosity | Fluid density |
| Unit | mm | mm | Mm | $\mu\text{L}$ | Pa | ° | rpm | mPa *s | kg/m<sup>3</sup> |
| Value | 1.2 | 0.8 | 16.4 | 300 | 0/40 | 10-20 | 2-10 | 1 | 1001 |

### 3) Endothelial lining and sprouting on-chip

Vasculature provides oxygen and nutrients for most tissue and supports organ function and physiology. The lining of the luminal surface of the vasculature depends on the substrate as well as on the mechanical features, also called haemodynamic forces, such as flow, pressure, speed and direction that translate into shear stress on the EC membrane. Hence, WSS is a key parameter influencing the morphology and physiology of ECs as demonstrated in different studies^22,23^.

To evaluate the behaviour of ECs in the rOoC platform, we first tested the viability and morphology of human umbilical vein endothelial cells (HUVECs) cultured over 2 weeks in the inner and outer PCs. For this, the OCs were filled with ECM (Geltrex), and subsequently, both outer and inner PCs of the rOoC chip were coated with a mixture of collagen I type and Geltrex in a 1:1 ratio. HUVECs were plated in a concentration of 2 × 10^6^ cells/mL and left to attach for 1 h without tilting. Then, perfusion was performed by placing the rOoC on the 3D-tilting platform. HUVECs formed a confluent monolayer covering the walls and the bottom of the PCs 24 h after plating. The confluent monolayer was preserved for at least 14 days of culture under both high and low WSS (Suppl. Fig 1). Elongation and alignment of ECs are the main morphological changes that can be observed within hours of exposure to flow stimulation^24–26^. Previously, it has been demonstrated that the alignment of ECs *in vivo* is characteristic of large blood vessels with uniaxial and laminar flow. Correspondingly, the ECs alignment *in vitro* is stronger for a uni-directional flow compared to a bi-directional flow^27^. We used the following settings to investigate alignment and sprouting (WSS values from Fig. 2d/e for inner/outer PC):

The cells were cultured for 4 days in the outer and inner PCs using the high-WSS setting, fixed and subsequently stained with FITC-conjugated phalloidin and DAPI to visualize the actin cytoskeleton and the nuclei respectively. Representative confocal images in Fig. 3a illustrate a clear difference between cells cultured under static conditions, and directional flow. This is not only apparent in the alignment of the actin fibres as shown in the magnified images (Fig. 3a, right side), but also in the alignment of nuclei (Fig. 3b). To quantify HUVECs alignment to the flow direction, orientation histograms were created for both actin filaments (Fig. 3a) and nuclei (Fig. 3b). The analysis was expanded to different flow settings, and a comparison between inner and outer circuits (fluorescence images and alignment histograms Suppl. Fig. 2). Obtained results demonstrated stronger alignment in the outer PC compared to the inner PC and for increased tilting speeds (high-WSS vs medium-WSS). This corresponds with the WSS values for both PCs and settings given in table 2.

**Table 2:** Flow settings and WSS values used in this study for the inner and outer PC

| Setting | Tilt [$^\circ$] | Speed [rpm] | WSS<sub>inner</sub> [dyne/cm<sup>2</sup>] | WSS<sub>outer</sub> [dyne/cm<sup>2</sup>] |
| --- | --- | --- | --- | --- |
| high-WSS | 15 | 2 | 1.5 | 3 |
| medium-WSS | 15 | 1 | 0.8 | 1.5 |
| low-WSS | 10 | 1 | 0.3 | 1.2 |

**Figure 3.**
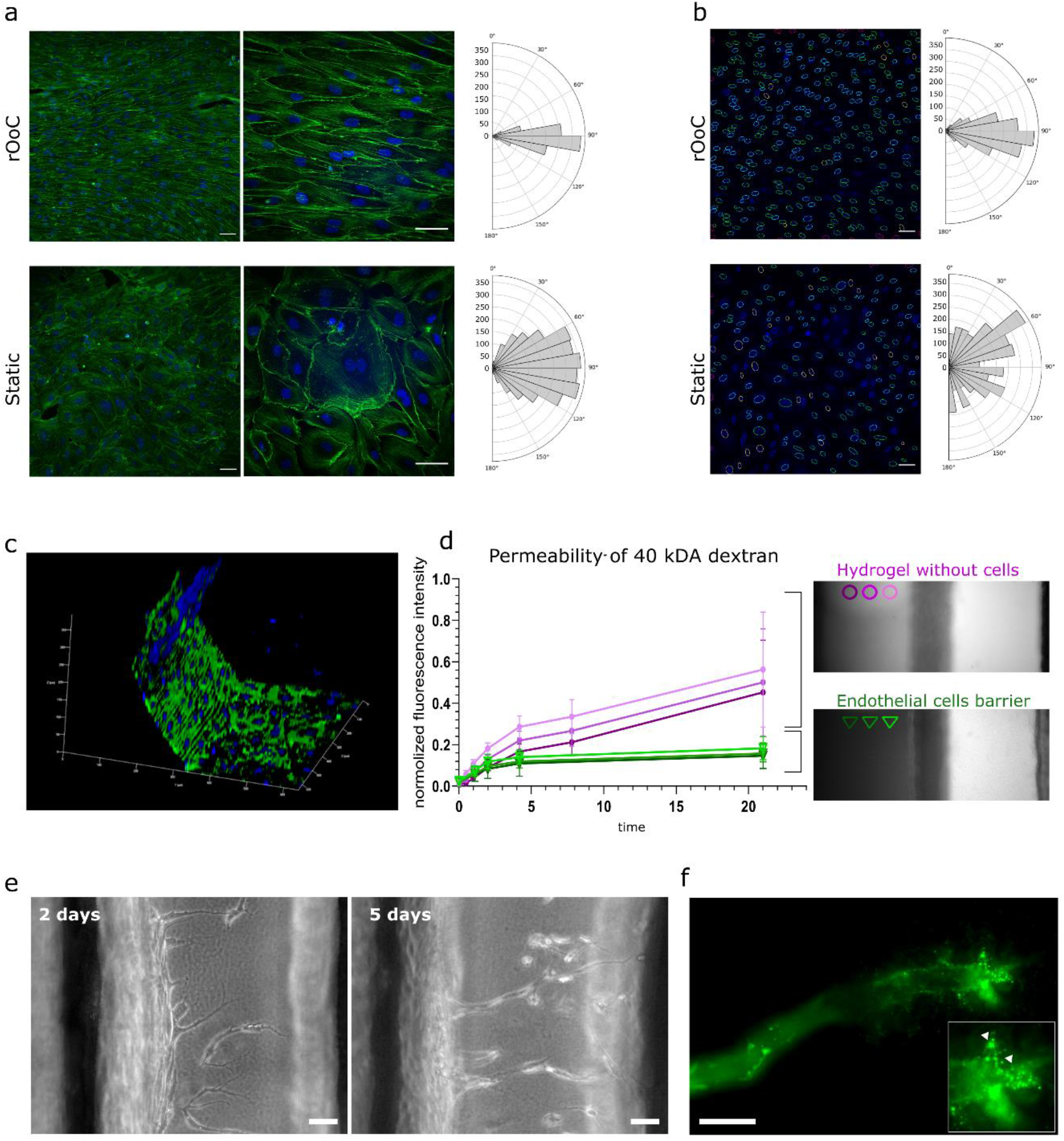
rOoC support alignment, barrier and vessel formation of human ECs. a) EC alignment and corresponding reorganization of actin cytoskeleton exposed to flow in the outer PC of the rOoC. Immunofluorescent staining of Actin (green) and nuclei (blue) of cell populations in the channel centre. Histograms representing vector counts in the area are calculated as a sum for three different images. Scale bar 50 μm. b) Alignment of HUVECs nuclei with the flow. Histograms representing nuclei counts in the area are calculated as a sum for three different images. c) 3D reconstructed confocal image of the endothelial barrier at the interface with the OC (blue – nuclei, green – VE-cadherin). d) Permeability test at the interface between the OC and the outer PC without cells and covered with HUVECs. Left: graph represents changes of the ratio of the averaged fluorescence intensity in 3 regions in gel vs outer PC, error bars show standard deviation (n = 4), magenta - outer PC without cells, green - outer PC covered with HUVECs. Right: Dextran distribution in the OC without (magenta) and with an endothelial barrier (green) at 21 h after introduction of 40 kDa FITC-Dextran into the outer PC (details in Appendix A). e) Angiogenesis assay in rOoC: representative bright field images show the appearance of tip cells after 48h of culture, and formation of the lumen by stalk cells after 5 days of culture. Scale bar 50 μm. f) Fluorescence image showing FITC-labelled microbeads perfusion in the lumen of a newly formed microvessel. Magnification shows a magnification of the tip of the microvessel (arrowheads mark FITC-labelled microbeads). Scale bar 50 μm.

Next, we evaluated the formation of an endothelial barrier in the rOoC. As mentioned above, ECs formed a confluent monolayer not only on the bottom but also on the walls of the PCs (Fig. 3c). The barrier formation in the outer channel was evaluated at the interface with hydrogel by the ratio of fluorescence between the gel region and the outer PC (Fig. 3d). For this, EGM2 medium was supplemented with 40 kDa FITC-conjugated dextran (250 μg/ml) and introduced into the outer PC with or without HUVECs similarly to the experiment done in Appendix A. The permeability assay confirmed the establishment of an effective barrier preventing dextran leakage (Fig. 3d).

The sprouting of new microvessels from a monolayer of ECs defines the initiation of angiogenesis and is an important process in physiological and pathological cascades. To test ECs sprouting on the rOoC, we plated HUVECs in the outer PC as described above and performed perfusion at the low-WSS setting (table 2). This setting has been empirically optimised and defined as an optimal setting for sprouting. The first tip cells in the ECM were observed after 48h of culture in the HUVEC endothelialized rOoC. To stabilize newly formed microvessels and promote their growth, we generated a gradient of angiogenic factors in the ECM compartment, as described in the methods part. From day 5 onwards, we observed the formation of microvessels with discernible lumens that extended towards the inner PC. Using GFP-labelled microbeads that were added to the endothelialized PC on day 6 after sprouting, we demonstrated that the newly formed microvessels are perfusable as beads were detected at the tip of these microvessels (Fig. 3e).

### 4) Liver organoids on-chip

Next, we tested whether the rOoC platform supports the viability and functionality of 3D liver organoids cultured in the OC. For this, we differentiated human embryonic stem cells (H1 line) and induced pluripotent stem cells (WTC-11 and WTSIi013-A cell lines) towards hepatocyte-like cells grown as 3D organoids (ESC-HLC, iHLC_1 and iHLC_2, correspondingly) together termed as HLC organoids. Protocol and characterization of the liver organoids are shown in Suppl. Fig 4. After 24 days of differentiation, HLC organoids were mixed with Geltrex in a 1:2 ratio and introduced into the OCs (5-20 organoids each). HLC organoids were cultured in the rOoC platform for up to 2 weeks.

**Figure 4.**
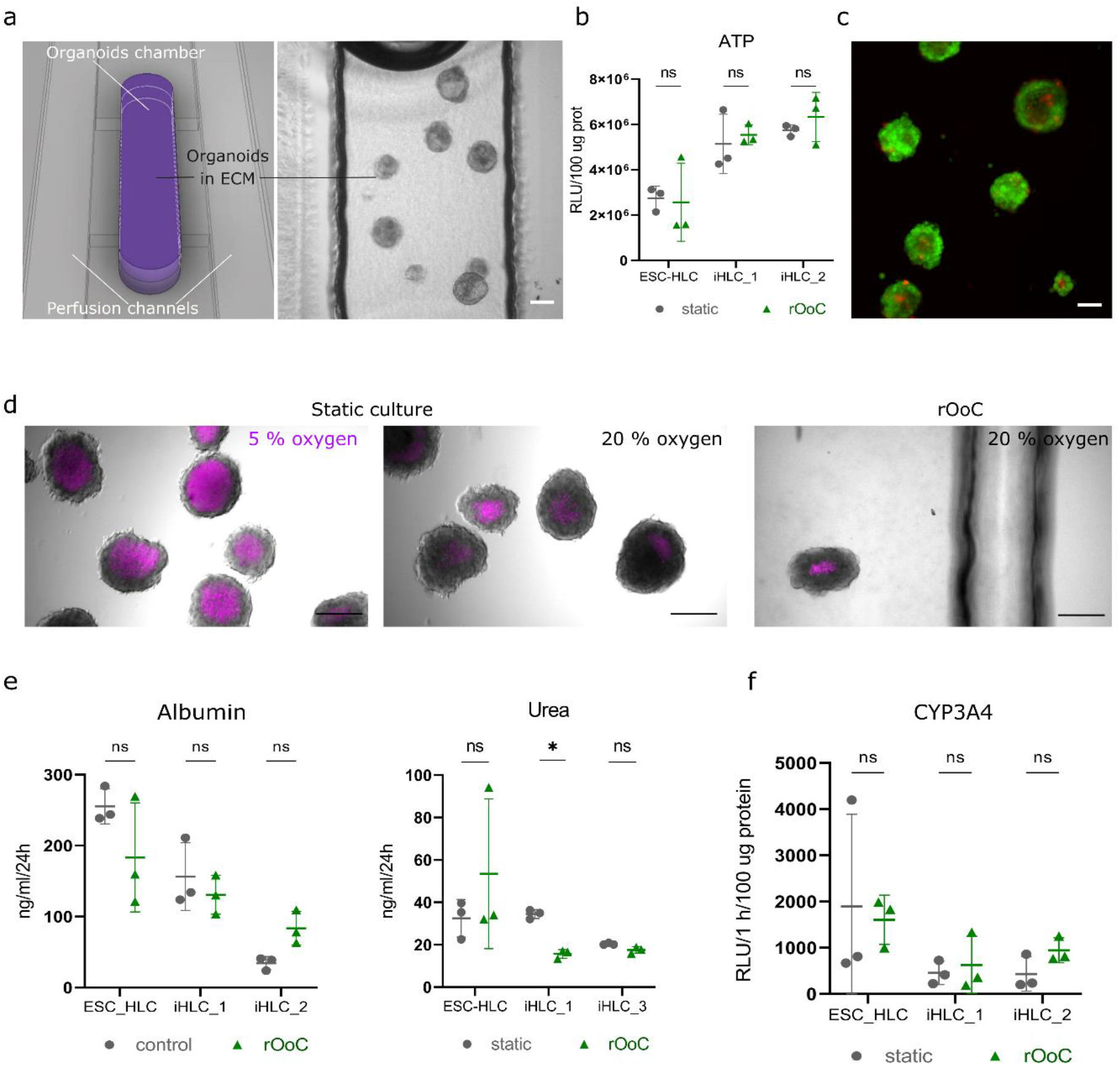
rOoC provides a platform for culturing 3D HLC organoids. a) 3D model of the OC in the rOoC platform and representative bright field image of iHLC organoids in the organoid culture chamber. b) ATP content of HLC organoids cultured for 2 weeks under static conditions and in the rOoC platform. c) Representative LIVE/DEAD image of iHLC_1 organoids in the rOoC. d) Image-iT™ Red Hypoxia Reagent staining of iHLC_1organoids under static conditions under hypoxia (5% (v/v) oxygen) and normoxia (20% (v/v) oxygen), and in rOoC under normoxia. e) Albumin and urea secretion of HLC organoids from 3 cell lines (n = 3 replicates), cultured in static conditions and the rOoC. The significance was calculated using a multiple paired t-test, *p < 0.05.) CYP3A4 activity of HLC organoids from 3 cell lines (n = 3 replicates) was evaluated by Luceferin-IPA (relative luminescence units, RLU, normalized to protein content) in static conditions and on the rOoC.

During this time HLC organoids preserved their overall morphological characteristics (Fig. 4a), however, some organoids exhibited sprouting of non-hepatic cells, due to the presence of an endothelial lineage in the HLC organoids. After 2 weeks, organoids in the rOoC had the same level of viability as in standard 3D static cell culture, without significant changes in ATP content (N=3 PSC lines) (Fig. 4b). Live/dead staining after 2 weeks of culture was also performed to visualize cell viability in the HLC organoids (Fig. 4c). To further characterize the culture conditions in the OC, we stained the HLC organoids after 48h of on-chip culture in the standard incubator (20% (v/v) oxygen, 5% (v/v) CO_2_, 37°C) with Image-iT™ Red Hypoxia Reagent (ThermoFisher Scientific, Fig. 4d). As a static positive control, we used free-floating organoids cultured in a hypoxia incubator (5% (v/v) oxygen), and as static negative controls free-floating HLC organoids placed in a standard incubator. Unlike the positive controls, organoids cultured on-chip showed a similar number of hypoxic cells as the negative control, mostly restricted to the organoids core as commonly seen. However, when more than 20 organoids were cultured in each OC, HLC organoids experienced hypoxic conditions, especially in the centre of the OC indicating a reduction of oxygen supply in these areas.

To test the functionality of HLC organoids, we compared the albumin and urea production of HLCs cultured under static conditions in an ultra-low attachment plate, or cultured in the rOoC (Fig. 4e). HLC organoids in the rOoC exhibited a similar level of albumin and urea secretions as those cultured under static conditions. Next, we tested CYP3A4 enzymatic activity, essential for the biotransformation of more than 30% of all prescribed drugs in the liver, in HLC using a luciferase-based assay based on Luciferin IPA (Promega, Sweden). The luciferase-based assay (Fig. 4f) indicated the same level of CYP3A4 activity in HLC organoids cultured in the rOoC platform and under static conditions.

Taken together, we demonstrated that the rOoC is suitable for the culture of 3D organoids, taking the example of HLC organoids, and supports their viability and functionality over at least 2 weeks of culture.

### 5) Circulating immune cells

Immune cells, both tissue-resident and circulating, play a fundamental role in nearly all diseases. While tissue-resident immune cells may be included in an organoid, immune cell circulation and homing are more challenging to reproduce *in vitro,* requiring either a Transwell® or a microfluidic setting^28^. Maintaining circulating immune cells in a microfluidic system can be difficult due to immune cell trapping and activation. Trapping may occur in tubing, reservoirs and internal pumps, reducing the number of cells that can interact with stationary tissue models. Unwanted activation of immune cells may occur due to mechanical stress^29^*e.g.* in areas with high shear forces or, in the case of internal pumps, by moving parts. Activation of immune cells changes the characteristics of the cells and may superimpose an effect that is alien to the intended physiological or disease model.

To determine viability as well as potential trapping and activation of immune cells in the rOoC platform, we isolated peripheral blood mononuclear cells (PBMCs) from 3 healthy donors as described in the methods part. Next, PBMCs were introduced in the outer PC of the rOoC system at a density of 3 × 10^5^ cells/ml in RPMI medium and the device was perfused on the platform to produce the previously defined high-WSS condition (table 2) in the outer PCs. The movement of the PBMCs on-chip was tracked using a miniaturized digital microscope (Dino-Lite Edge 3.0, Dino-Lite Europe B.V., Netherlands). As shown in the Suppl. Video 2, PBMCs followed the general flow in the PCs without any visible trapping at any point of the chip. Next, we cultured the PBMCs for 24h either under static or flow conditions followed by staining the cells with a non-viability marker (7AAD). Next cells were labelled with monoclonal antibodies to determine cellular subsets and possible activation by flow cytometry. As seen in Fig. 5, PBMCs that were maintained on-chip for 24h did not have a significant decrease in cell viability (97,05 ± 2,58% in static culture vs 90,42 ± 6,41% in rOoC, p=0,093) nor did they show significant changes in the percentages of T-cells populations with early (CD69) or late (HLA-DR) activation markers.

**Figure 5.**
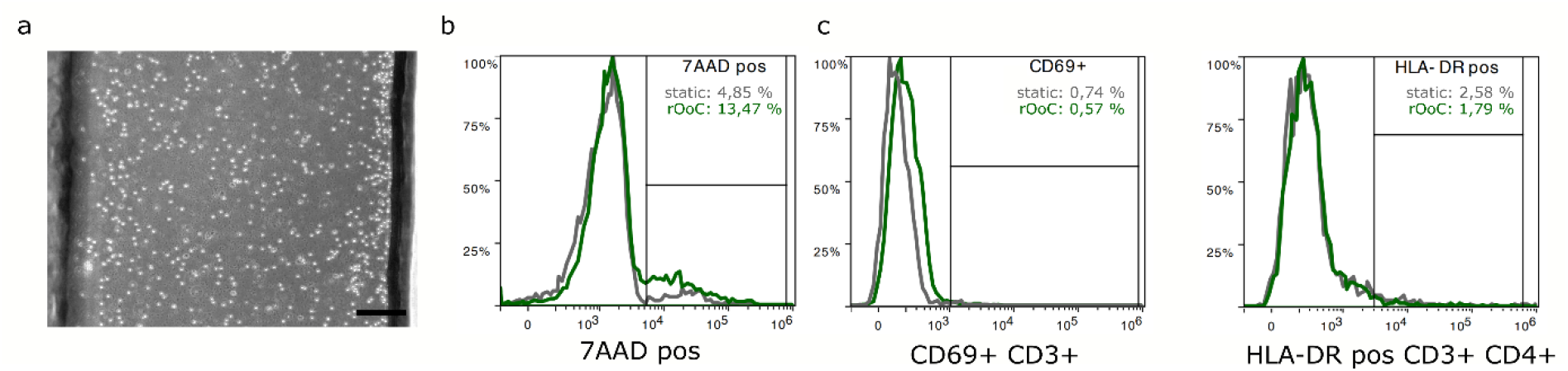
rOoC maintains recirculating PBMC phenotypes and viability. a) Homogeneous distribution of PMBCs in perfusion channel of the rOoC. Representative bright field images, scale bar 200 m. b) Representative histograms showing the percentage of non-viable (7AAd positive) cells in static culture (grey) and in the rOoC (green). c) Representative histograms showing the percentage of T cells (CD3+) with the early (CD69) activation marker and T-helper cells with the late (HLA-DR) activation marker in static culture (grey) and in the rOoC (green) after 24 h.

## Discussion

We present a novel, pump-less rOoC platform capable to cultivate different types of cells or organ representations in a closed microfluidic circuit with a uni-directional flow. The device is built of laser-structured thermoplastic films that are thermally bonded to form a multi-layer device. One of the main advantages of this technology over the often used soft lithography process^30^ is that thermoplastic films do not adsorb small molecules to the extent to which soft lithography materials do (especially Polydimethylsiloxane (PDMS)^31^).

Besides fabrication, our layout presents several advantages over pump-driven platforms: air bubbles are automatically trapped in the reservoirs avoiding channels blockages and cells death, cell/organoid loading and media exchange are simply done by pipetting and many systems can be run in parallel (8 rOoCs per plate, >10 plates per rotator). Compared to pump-driven systems that require controlling units and connecting tubes, the rOoC has higher scalability and lower cost^32^ and does not require the integration of flexible membranes^17^ for fluid manipulation. The integration of flexible membranes increases fabrication time and costs and potentially influences the behaviour of biological material by the introduction of another material into the system and by the implementation of local high pressure which will in particular affect circulating cells. Compared to other gravity-flow-driven systems, the rOoC generates a uni-directional instead of a bi-directional flow. This results in a better alignment of ECs (Fig. 3) and allows for the interaction of different organ representations^27^.

As organoids are the main building block in 3D cell culture, we demonstrated the suitability to embed and culture liver organoids in the rOoC. For this, we use a specific chamber layout with a stepped channel geometry that can be used to structure the ECM and provide a biologically relevant and biodegradable separation between PCs and OCs^21^. We have shown the organoid functionality using stem-cell-derived liver organoids and proved that the organoids stay healthy, not showing a significant hypoxic core, secrete albumin and urea and show drug metabolizing activities at the same level as the static control.

The advantage of an ECM-based separation layer instead of a separating polymer membrane is that this layout can be used to vascularize tissue representations. We have demonstrated that the platform allows EC to sprout into the OC. To achieve long-term stable perfusable microvessels, an interstitial flow^33,34^ will be added in a future iteration of the device. To do so, the layout can be adapted to generate a sufficient pressure gradient between both perfusion channels, *e.g.* by using bigger reservoirs or placing both perfusion circuits next to each other instead of one in the other. This will lead to an alternating and not parallel flow direction in both PCs thus generating a higher pressure gradient as with the current configuration.

Finally, we showed that circulating immune cells can be cultured without trapping in external tubing, integrated pumps or accumulation in medium reservoirs. At the same time, the ECM separation layer allows for immune interaction with the tissue representations cultured in the OC e.g. to study immune cell homing or inflammatory processes.

Most importantly, we have thoroughly characterized the rOoC and developed a computational model to predict flow rates and WSS values. This will help us to conduct a model-driven design approach in the future to tune the platform to new biological questions. The flow characteristics of a fluidic system can be classified using the Reynolds (Re) and the Womersley (Wo) numbers (Definition see Suppl. Text). Re relates to the relationship between viscous to inertial forces in the liquid flow, whereby high values of Re lead to turbulences. Wo is used to describe if inertial effects due to unsteady pressure lead to a disturbed flow profile (oscillatory flow). The maximum Re number in the rOoC is 115, which is clearly within a laminar regime and the maximum Wormesley number in our system is Wo_max_ = 0.2 thus the flow can be considered undisturbed by transient effects (Calculations and channel dimensions see Suppl. Text). Given these flow conditions, we mimic healthy conditions for EC lining^24^. The flow rate/WSS can be tuned either by design or by changing the driving conditions (tilt, speed, volume) thus we can model an arterial and a venous side (different WSS) in one device. For example, as described in Appendix C, a high reservoir volume will result in a bi-directional flow pattern that is associated with a bi-directional flow pattern resulting in loss of EC formation and is associated with inflammation and heart diseases^24^.

In summary, the rOoC platform provides a robust and scalable new tool for building comprehensive *in vitro* models by incorporating three central components: long-term cultures of functional endothelial cells, ECM-embedded 3D organ representations and circulating immune cells. This, together with the versatility of the flow regulation should provide a valuable next-generation platform for disease modelling and interrogation.

## Materials and methods

### Device fabrication

The rOoC chips are made of several laser-machinedPMMA sheets (Hesaglas®, Topacryl AG, Switzerland) thermally bonded together. First, each layer was designed in Autodesk Inventor® 2019 (Autodesk Inc., USA) and saved as individual DXF files for laser machining. The top part (6 mm thick) containing the reservoirs was processed with a conventional CO_2_ laser cutter (Beambox®, Flux Inc., Taiwan). Reservoirs were laser-engraved using a power of 12W and a speed of 25 mm/s whereas holes and the outline was cut with a power of 24W and a speed of 6 mm/s. Thin PMMA films (0.3/0.5 mm) containing the PCs and OCs are laser micro-structured using an RDX-500(Pulsar Photonics GmbH, Germany) equipped with a UV (343 nm) femtosecond laser (Pharos PH2-10 W, Light Conversion, Lithuania). The films were cut with a speed of 500 mm/s and a peak fluence of 11.03 J/cm^2^.

After structuring, the films were cleaned in an ultrasonic bath (USC100T, VWR International, USA) for 15 min, sprayed with a nitrogen gun, and dried in an oven at 65 °C for 15 min. Next, the substrates were UV-activated using an Excimer lamp with a 172 nm emission wavelength (ExciJet172 55-130, Ushio GmbH, Germany). The distance to the lamp was adjusted to 1 mm and the substrates were exposed to a dose of 0.6 J/cm^2^. Finally, the parts were aligned on an aluminium plate using alignment pins and transferred to a pneumatic hot-press (AirPress-0302, Across International LLC, USA). For equal pressure distribution and control, cleanroom paper was put on top of the stack and an electronic pressure regulator (ITV1030, SMC Corporation, Japan) was used to control the bonding pressure. After UV activation, PMMA can be thermally bonded below its glass transition temperature (T_g_ = 100 °C). We used a bonding pressure of 2.4 MPa, a temperature of 84 °C, and a bonding time of 15 min. In a previous study, test devices were fabricated with this technology that withstood loads >40 N with a thermal-induced channel collapse of less than 6 μm for a 200 mm^2^ large chamber.^17^

To prevent free-standing structures (e.g. in layer 2 or 3) from falling apart, small bridges (0.5 mm wide) were included that were later removed using a knife. This made it necessary to first bond the top parts of the device separately and later seal them in a second step (see Fig. 5). For on-chip confocal imaging, the devices were sealed with a 175-μm thick PMMA film (Merck Sigma, USA).

**Figure 5:**
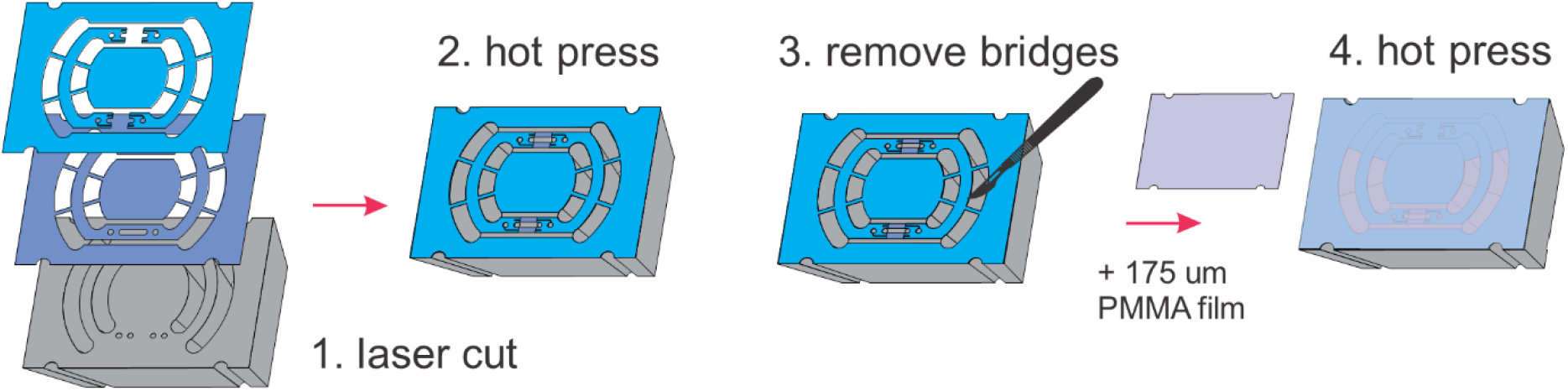
Fabrication technology using laser-cut PMMA films thermally bonded together

### Flow measurement

The flow was measured by micro-Particle Image Velocimetry with a setup as described earlier^35^. As tracers, dyed 10-μm polystyrene microparticles (Merck Sigma, USA) in a concentration between 0.1% and 1% (m/v) were used. Images were acquired in transmission by using a digital microscope (H800-96S-ES3, United Scope LLC, USA) coupled with a high-speed camera (acA1300-200uc, Basler AG, Germany). The microscope was mounted to a customized stand including a LED backlight illumination as shown in Suppl. Fig. 5. The stand was assembled using laser-cut PMMA sheets and mounted on the moving platform. Particle movement was observed in a field of view of 592 × 100 Px (1 Px = 5.33 μm) at 2270 fps and 50 μs acquisition time. Later, image stacks of 10.000 frames were evaluated using an optical flow algorithm previously applied to calculate the motion of beating cardiomyocytes^36^. The setup is suitable to detect flow velocities up to 100 mm/s. The measurement was calibrated by applying defined flow rates to channels with a similar cross-section using a syringe pump as a pressure source (Aladdin AL-1000, World Precision Instruments LLC, USA). A calibration factor of 1.84 ± 0.4 was determined to calculate the flow rate Q from the measured velocity v_m_ and the channel cross section (b, w):

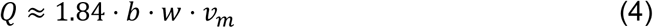

Moreover, the measurement was validated using the transported volume (*V*(*t*) = ∫ *Q*(*t*)). This volume needs to be the same as the amount of liquid filled in the reservoir. The Suppl. Table shows the flow data used for validating the model and the respective volume. Finally, due to the unsteady nature of the flow, it has to be ensured that the flow is precisely resolved with the setup. One important aspect is the type of tracers used and whether they follow abrupt changes in the flow velocity. A way to quantify this is by calculating the Stokes number^37^ St or the particle relaxation time τ. It can be calculated given the particle density 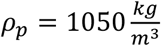, the particle diameter *d*_*p*_ = 10*μm*, and the viscosity from Table 1:

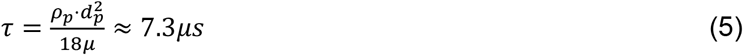

That means, the tracers need roughly 40 μs after a sudden change in the velocity to adapt to the new velocity (tracing error < 0.5% for a decay function of 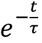). This value is much lower than the recording frequency of 2270 fps thus it can be stated that the particles follow the flow to a good extent.

### Computational model

The flow in the rOoC is driven by the pressure difference between the inlet and outlet:

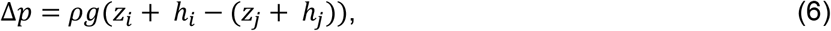

where *z* is the current elevation of the PC at the inlet/outlet and ℎ is the orthogonal distance to the liquid water table, g is the gravity constant (g = 9.81 m/s^2^) and *ρ* is the fluid density. We assume a fully developed laminar flow in the PCs. The transmissibility *σ* of the channel is obtained analytically for straight rectangular channels, or by solving the Navier-Stokes equations for more complex channel geometries. The flow rate in a rectangular channel (length: l_c_, width: b, height: w) is approximated by the analytical solution for fully-developed laminar flow in a rectangular duct (Suppl. Text). Forming the volume balance of the reservoirs yields:

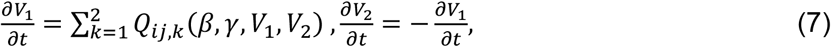

where the flow rate function *Q*_*ij*,*k*_ depends on the volumes in both reservoirs and the pitch and roll angles (via the elevation and water table-dependent pressure difference). Moreover, the volume in both reservoirs is always greater than zero.

Based on these equations, we first developed a simplified model to understand the basic physics involved in the system (Appendix C). As it turned out, several additional effects needed to be taken into account:

- Due to the curved geometry of the reservoir, the water table height ℎ non-linearly depends on the water volume and current tilt angles. To determine the liquid level in the reservoirs, we intersect the tilted reservoir geometry with the horizontal plane such that the intersected volume equals the current filling volume (determined with Eq. 7). This nonlinear problem is solved iteratively using Brent’s method.
- Since the reservoirs have a fixed maximum volume (V_max_), the volume balance ensures that the volume never exceeds this value.
- The abrupt dimension change at the in-/outlets of the reservoirs acts as capillary-stop-valves^38^. If the reservoirs run dry at one in-/outlet, the air-liquid interface will prevent backflow until a critical pressure p_c_ is reached. This pressure is defined by the surface tension δ of the medium, the contact angle on the plastic surface θ and the estimated curvature radius of the liquid-air interface R:

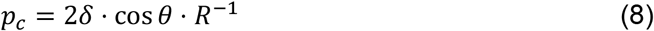 We measured the respective contact angles and fabricated a microfluidic test device to determine the critical pressure experimentally (see Appendix C).

### Endothelial cells culture

Human vein endothelial cells (HUVECs, Life Technologies, catalogue no. C0035C) and human liver endothelial cells (HLECs, Lonza, catalogue no. HLECP2) were routinely cultured in T75 flasks (Thermo Fisher Scientific, catalogue no. 156499) with Endothelial Cell Growth Medium EGM2 (Lonza, catalogue no. CC-3162). Cells between passages 3 and 5 were used for the experiments. Media was replaced every 2 days. All cell cultures were tested negative for mycoplasma.

Before cell seeding into the rOoC, all devices were sterilized by rinsing with 60% (v/v) ethanol and water and subsequently dried under UV. A mixture of Geltrex (ThermoFisher Scientific, catalogue no. A1413302) and Collagen type I (ThermoFisher Scientific, catalogue no. A1048301) in ratio 1:1 was used as a 3D scaffold (bellow referred to as the extracellular matrix, ECM). A stock solution of 3 mg/mL rat tail collagen type I was neutralized with 1% 1 M NaOH (Sigma, S5761) and 10% 10x phosphate buffered saline, PBS (Thermo Fisher Scientific). The neutralized collagen was kept on ice until use and used within 10 min. The organ chamber was filled with 17 μL of ECM. After polymerization of ECM (incubated for 10 min at 37°C), PCs were coated with 1% (v/v) of the same ECM diluted in a cold DMEM/F12 medium. Endothelial cells were dissociated by incubation with Trypsin/EDTA (Sigma, T3924-100ML), and re-suspended in EGM2-medium at a concentration of 2 × 10^6^ cells/mL. 50 μL of the cell suspension was dispensed into the PCs and incubated for 1 h at 37 °C, with 5% CO2. For an even distribution of the cells to both walls, we first placed the rOoC on the left side and let the cells attach for 45 to 60 min. Next, a second aliquot of cell suspension was added to the PCs and the rOoC was placed on the right side for the same time. After the cells were attached to both walls, the rOoC was incubated for another 45 min to let cells attach to the bottom of the PCs. After that rOoC was placed on the 3D tilting platform. The following settings of the platform were used in the experiments with endothelial cells: speed 1-5 rpm, and tilt of 10-17°. The medium was changed daily.

### Endothelial cells sprouting assay

HUVECs were cultured for 48 h in the outer PC of the rOoC using a speed of 2 rpm and a tilt of 15°. After observation of the first tip cells, the EGM2 medium in the inner PC was supplemented with three angiogenic factors: vascular endothelial growth factor A (VEGF-A) at 20 ng/ml, Sphingosine-1-Phosphate (S1P, Sigma, S9666) at 500 nM and phorbol myristate acetate (PMA) (Sigma, P1585) at 20 ng/mL. Sprouting was observed for 7 more days.

### Perfusion with beads

To check the perfusability of formed vessels EGM2 medium was supplemented with FluoSpheres™ Polystyrene Microspheres, 1 μm (ThermoFisher Scientific, catalogue no. F13080) introduced in the outer PC with endothelial cells. The presence of fluorescent particles in the microvessels was evaluated after 1h of perfusion using a fluorescent microscope (Zeiss, Axioscope) with a 10x objective.

### Endothelial permeability visualization

After 4 and 7 days of culture, EGM2 supplemented with 10 and 40 kDa FITC-conjugated Dextran (Sigma, FD40-100MG) in a concentration of 250 μg/mL was added to the outer PC, covered with cells. The permeability was evaluated at the interphase with ECM. Time-lapse images were acquired at 60 min intervals using a fluorescent microscope (Zeiss, Axioscope) with a 10x objective. Fluorescence intensity in the outer PC (donor site) and ECM (recipient site) was analyzed using Fiji software.

### Generation of hepatocyte-like cell organoids

Human embryonic stem cells (H1, Coriell Institute for Medical Research) and human induced pluripotent stem cells (HLC_1: WTC-11, Coriell Institute for Medical Research; HLC_2: WTSIi013-A Wellcome Trust Sanger Institute) were routinely cultured in mTeSR media (StemCell Technologies, catalogue no. 85857) human E8 media (Thermo Fisher Scientific, catalogue no. A1517001) correspondingly, on plates coated with 0.1% (v/v) Geltrex (Thermo Fisher Scientific, catalogue no. A1413201) in a humidified 37 °C, 5% CO_2_ incubator. The pluripotency of cells before differentiation was confirmed by flow cytometry, qPCR, and immunofluorescent imaging for pluripotency markers. Normal karyotype and absence of mycoplasma contamination were confirmed for all used cell lines.

Hepatic differentiation was performed using a modification of previously published protocols^39^. Briefly, iPSC/ESCs were differentiated toward definitive endoderm in IMDM/F12 media containing 1% (v/v) lipid concentrate (Thermo Fisher Scientific, catalogue no. 11905031), 100 μg/ml transferrin, 3 μM CHIR99021 (Tocris Bioscience, catalogue no. 4423), 50 nM PI-103 (Tocris Bioscience, catalogue no. 2930) and 100 ng/ml activin A (Peprotech, catalogue no. 120-14P) for 24 h and 100 ng/ml activin A for subsequent 48 h. The definitive endoderm cells were treated with 10 ng/mL FGF2 (Peprotech, catalogue no. 100-18B) and 20 ng/mL BMP4 (Peprotech, catalogue no. 120-05) in IMDM/F12 medium supplemented with 1% (v/v) N-2 (Thermo Fisher Scientific, catalogue no. 17502-048), 1% (v/v) B-27 minus vitamin A (Thermo Fisher Scientific, catalogue no. 12587010) and 1% (v/v) lipid concentrate, then with 5 μM A8301 (Stem Cell Technologies, catalogue no. 72022), 20 ng/mL HGF (Peprotech, catalogue no. 100-39H), 20 ng/mL BMP4, 1% (v/v) B-27 with vitamin A for 3 more days and with 25 ng/mL HGF, 1% (v/v) DMSO for another 5 days. At day 12, cells were detached by accutase and aggregated in the agarose U bottom microwells in the presence of 25 ng/mL HGF, 0.1 μM Dexamethasone, 10 μM Y-27632, 0.5% (v/v) ITS, 0.1% (v/v) lipids concentrate, 100 μM Ascorbic acid-2 phosphate (AAP), 1% (v/v) B-27 (without vitamin A) and 1% (v/v) N-2. After formation of spheroids at day 13, media was replaced with William’s E media, supplemented with 5% (v/v) FBS, 20 ng/ml HGF and 10 ng/ml oncostatin M (Peprotech, catalogue no. 300-10), 1% (v/v) ITS, 100 μM AAP, 0.1 μM Dexamethasone. For further maturation, organoids were cultured in microwells in William's E media, supplemented with 1% (v/v) ITS, 0.1 μM Dexamethasone, 20 ng/ml Oncostatin M and 1% (v/v) MEM Non-Essential Amino Acids Solution (Thermo Fisher Scientific, catalogue no. 11140050), 10 uM DAPT for another 10 days with the replacement of media every 48h. On day 18, organoids were additionally incubated for 1 h in the same medium supplemented with 5% (v/v) of Geltrex.

### Hypoxia test

Visualization of hypoxia in HLC organoids was performed using Image-iT™ Red Hypoxia Reagent at a concentration of 5 μM according to the manufacturer’s instructions. Organoids were imaged using a fluorescence microscope (Zeiss, Axiovert) with excitation/emission of 490/610 nm and 10x objective.

### Albumin and Urea production assays

Albumin content in the supernatant media was evaluated using Human Albumin ELISA Quantitation Set (Bethyl Laboratories, catalogue no. E88-129). The concentration of urea in the medium was quantified calorimetrically using a urea assay kit (BioAssay Systems, DIUR-100), following the manufacturer’s instructions. Both albumin and urea concentration was normalized to the total protein content, using Pierce™ BCA Protein Assay Kit (Thermo Fisher Scientific) according to the vendor's instruction.

### Viability and hepatotoxicity

ATP content was evaluated using Cell Titer-Glo® 3D Cell Viability Assay (Promega, Sweden) according to the manufacturer’s instructions.

### Cytochrome CYP3A4 activity

Cytochrome CYP3A4 enzymatic activity of HLCs was measured using P450-Glo™ Assay with Luciferin-IPA (catalogue no. V9001, Promega, Sweden). The relative luminescence was normalized to total protein content, measured by Pierce™ BCA Protein Assay Kit (Thermo Fisher Scientific) according to the vendor's instruction.

### Immunofluorescence staining and microscopy

Cell cultures in rOoC were fixed in 4% (w/v) PFA for 20 min on the orbital shaker. Each step was followed by three washing steps (each 10 min) in DPBS using an orbital shaking. Permeabilization and blocking was performed by incubation in PBS with 1% (m/v) BSA (Sigma Aldrich), 0.2% (v/v) Triton-X100 (Sigma Aldrich) and 0.5% (v/v) DMSO at RT for 2 h on the orbital shaker. Staining with primary antibodies was performed for 24 h (at 4 °C) with subsequently 2 h incubation with secondary antibodies (Jackson ImmunoResearch, West Grove, PA) diluted with 1 % BSA, 0.1 % Triton-X100 in PBS. Primary antibodies (Ab) used in this study: rabbit polyclonal Ab to human serum albumin (Abcam, catalogue no. ab2406, 1:400), goat polyclonal antibody to E-cadherin (R&D systems, catalogue no. AF748-SP, 1:250), mouse monoclonal Ab to CYP3A4 (3H8) (Invitrogen, catalogue no. MA5-17064, 1:250), rabbit polyclonal to VE-Cadherin (Abcam, catalogue no. ab33168, 1:400), mouse monoclonal to human CD31 (Abcam, catalogue no. ab24590, 1:200). Secondary Ab used in this study: Alexa Fluor® 488 AffiniPure Donkey Anti-Goat IgG (H+L) (catalogue no. 705-545-147, 1:300), Cy™3 AffiniPure Donkey Anti-Rabbit IgG (H+L) (catalogue no. 711-165-152, 1:400), Alexa Fluor® 647 AffiniPure Donkey Anti-Mouse IgG (H+L) (catalogue no. 715-605-150, 1:400) (all secondary Ab are from Jackson ImmunoResearch). Nuclear counterstaining was performed with 1 μg/mL Hoechst 33258 (Sigma Aldrich). Confocal microscopy was performed on a Zeiss 700 laser scanning confocal microscope using standard filter sets and laser lines with a 40x oil immersion objective. Images were acquired using Zen software (Zeiss) as Z-stacks with 2 μm spacing between stacks. The confocal images were analyzed using Fiji software^13^ and are displayed as a resulting Z-stack.

### RNA extraction and real-time polymerase chain reaction (PCR)

RNA was isolated using an RNeasy Micro kit (Qiagen, Germany) according to the manufacturer’s protocol. cDNA was synthesized using a High-Capacity cDNA Reverse Transcription Kit (Thermo Fisher Scientific, catalogue no. 4368814). Gene expression analysis was performed using a TaqMan Universal mix on a TaqMan ViiA7 Real-Time PCR System. The primers used in this study were: ALB (Hs00609411_m1), HNF4A (Hs00230853_m1), CYP3A4 (Hs00604506_m1), A1AT (SERPINA, Hs01097800_m1) and RPL (Hs02338565_gH), all purchased from ThermoFisher Scientific. RPL was used as endogenous control. The levels of expression of genes of interest were quantified by ddCt with normalization to undifferentiated stem cells (d 0) differentiated from the H1 line. Data represent three donor PHH samples, and HLC differentiated from 3 cell lines.

### PBMC isolation and culture

Human peripheral blood mononuclear cells (PBMC) were isolated by a density gradient technique, using the Lymphoprep™ gradient (StemCell Technologies, 07851) using the vendor’s protocol. PBMC were cultured for 24 h in an advanced RPMI medium (ThermoFisher Scientific, catalogue no. 12633012), supplemented with 2% (v/v) FBS, 1% (v/v) penicillin-streptomycin and then analyzed by FLOW cytometry.

### FLOW cytometry

PBMC were collected from the chips and centrifuged at 300g for 5 min. Cells were washed once with FACS buffer (PBS, supplemented with 1% BSA and 0.05% Sodium Azide) and centrifuged again at 300g for 5 min. Staining for immune subsets and activation markers was performed for 30 min, followed by a wash step with FACS buffer and analysis at a “Gallios” flow cytometer (Beckman Coulter Life Sciences, United States). The following antibodies were used at a dilution of 1:250 in FACS buffer:

- Anti-Human CD4 – FITC (eBioscience 11-0049-42), Clone RPA-T4
- Anti-Human CD3 – PE (BioLegend 300308), Clone HIT3a
- Anti-Human CD69 – PE/Cy7 (Biolegend 310912), Clone FN50
- Anti-Human HLA-DR – APC/Cy7 (Miltony Biotech 130-111-792), Clone REA805
- 7AAD ready-made solution (Sigma SML1633), dilution: 1:100

### Image processing

The alignment of ECs was analysed with two different toolchains for the cytoskeleton and the nuclei. Image pre-processing was done in FIJI^40^, cell/nuclei segmentation was performed with Cellpose^41^ and for post-processing, we used Python with the libraries Matplotlib and OpenCV. For actin-stained fluorescence/confocal and phase-contrast images the following process was applied:

a. For confocal images only: Z-projection in FIJI.
b. Conversion in an 8-bit image.
c. Analysis of cell alignment using the OrientationJ Vector Field^42^ plugin from FIJI:

- Gradient: Riesz-Filters
- Local window: 10 Px
- Grid size: 40 Px
d. The generated vector field is saved as a CSV file and loaded in a Python script (Suppl. Code 4) that extracts and plots the orientation histograms.

For the orientation of the nuclei the following toolchain was used:

I. For confocal images only: Z-projection in FIJI.
II. Conversion in 8-bit image and application of a Gaussian-blur filter (4 Px size).
III. Segmentation in Cellpose using the “nuclei method” and a size of 20 – 100 Px (depending on magnification) and saving as an ImageJ contour file.
IV. Reading the contour files in a Python script (Suppl. Code 5) that fits ellipses to the contours and calculates nuclei orientation and aspect ratio.
V. Plotting of nuclei orientation histograms using the script from d).

### Statistics

Statistical analyses and graph generation were performed using GraphPad PRISM 7 (GraphPad Software Inc.). Unless specifically stated, a two-tailed, paired t-test (with unequal variances) was applied for the comparison of the two groups. If not stated otherwise, a one-way ANOVA analysis was applied. The data are presented as mean ± SD. Statistical significance was assigned as not significant (NS) P > 0.05; or significant with *p ≤ 0.05; **p ≤ 0.01; ***p ≤ 0.001; ****p ≤ 0.0001.

## Supplementary Materials

**Table 3:** List of Suppl. material

| Name | Location | Description |
| --- | --- | --- |
| Suppl. Fig. 1 | Suppl-Fig-Tilting.pdf | Long-term culture of HUVECs on-chip |
| Suppl. Fig. 2 | Suppl-Fig-Tilting.pdf | Alignment of HUVECs in the inner/outer PC |
| Suppl. Fig. 3 | Suppl-Fig-Tilting.pdf | Culture of Human liver endothelial cells on-chip |
| Suppl. Fig. 4 | Suppl-Fig-Tilting.pdf | Organoid generation and differentiation |
| Suppl. Fig. 5 | Suppl-Fig-Tilting.pdf | Experimental setup for μPIV measurement |
| Suppl. Fig. 6 | Suppl-Fig-Tilting.pdf | Gating strategy for flow cytometry analysis |
| Suppl. Video 1 | animation-chip.gif | Modelled liquid volumes in the reservoirs |
| Suppl. Video 2 | Cells-tilt-15-2rpm.wmv | PBMCs flowing in the PCs |
| Suppl. Text | Channel_flow.pdf | Text describing the fluidic resistances of and characteristic flow numbers of the channels |
| Suppl. Table | Calibration-uPIV.xlsx | Flow data used to calibrate the μPIV setup |
| Suppl. Code 1 | https://github.com/MathiasBusek/Code-Tilting-Paper | Arduino code to measure the tilt of the platform using a gyroscope |
| Suppl. Code 2 | https://github.com/MathiasBusek/Code-Tilting-Paper | Python script used to generate plots for pitch and roll in Fig. 2b |
| Suppl. Code 3 | https://github.com/timokoch/dumux-microfluidics | Source code of the developed computational model based on DuMux<sup>43</sup> |
| Suppl. Code 4 | https://github.com/MathiasBusek/Code-Tilting-Paper | Python script used to calculate the orientation of the cytoskeleton in Fig. 3 |
| Suppl. Code 5 | https://github.com/MathiasBusek/Code-Tilting-Paper | Python script used to calculate the orientation of the segmented nuclei in Fig. 3 |
| Suppl. Code 6 | https://github.com/MathiasBusek/Code-Tilting-Paper | Python script used for the simplified flow model calculations in Appendix C |

## Author Contributions

M.B. developed the technology, designed and characterized the devices, prepared figures, and wrote the paper together with A.A. Furthermore, A.A. designed the biological part of the paper, performed cell experiments and prepared figures. T.K. developed the computational model and wrote the modelling part of the paper and the Suppl. Text. A.F., L.D., M.A.-M., C.D. and J.S. performed cell experiments and biological readouts. A.G. and S.G. fabricated the microdevices. S.K. and E.M. supervised the work, provided lab space, and procured funding for this study. All authors have read and agreed to the published version of the manuscript.

## Acknowledgement

Flow cytometry assays and quality control of iPSC lines were performed at The Norwegian Centre for Stem Cell Research, Oslo University Hospital.

This work received funding from the Research Council of Norway through its Centers of Excellence scheme; project No. 262613, open project support No. 315399 and a Qualification grant No. 329001. Furthermore, financial resources were granted from the South-Eastern Norway Regional Health Authority Innovation project No. 30629 and the European Union’s Horizon 2020 Research and Innovation program under the Marie Skłodowska-Curie Actions Grant, agreement No. 801133 (Scientia fellowship), PSC partners and the Norwegian PSC research centre.

## Conflicts of Interest

M.B., A.A., M.A.-M., and S.K. have applied for a patent covering the main principle of fluid actuation and application and are planning to commercialize the technology via a start-up company.

## Appendix A: Diffusion experiment

To test the capability of the rOoC device to establish and maintain gradients, the diffusion of fluorescein-labelled Dextran (10, 40 and 70 kDa, Merck Sigma, USA) from one side of the OC to the other was observed. First, the organ chamber was filled with an ECM (Geltrex, ThermoFisher Scientific, catalogue No. A1413201) to generate a flow barrier. Next, a dextran solution dissolved in PBS (concentration: 0,25mg/ml) was injected into the reservoirs either connected to the inner or outer PC (donor side). This setup is similar to the rOoC chip filled with red food colour in the inner circuit as shown in Fig. A1. Now, the fluorescence intensity in three different regions of interest (ROIs) (1, 2, and 3 marked in Fig. A2) was measured with a fluorescence microscope (Axio Vert. A1, Zeiss, Germany). Images were recorded every 5 min with a constant exposure time (28ms) and excitation intensity. Next, the mean intensity in the FOW was evaluated using FIJI^40^. Finally, the fluorescence signals were normalized to the maximum intensity and averaged over all ROIs. Fig. A3 shows the normalized fluorescence intensity for 70 kDa dextran depending on the distance from the PCs.

**Figure A:**
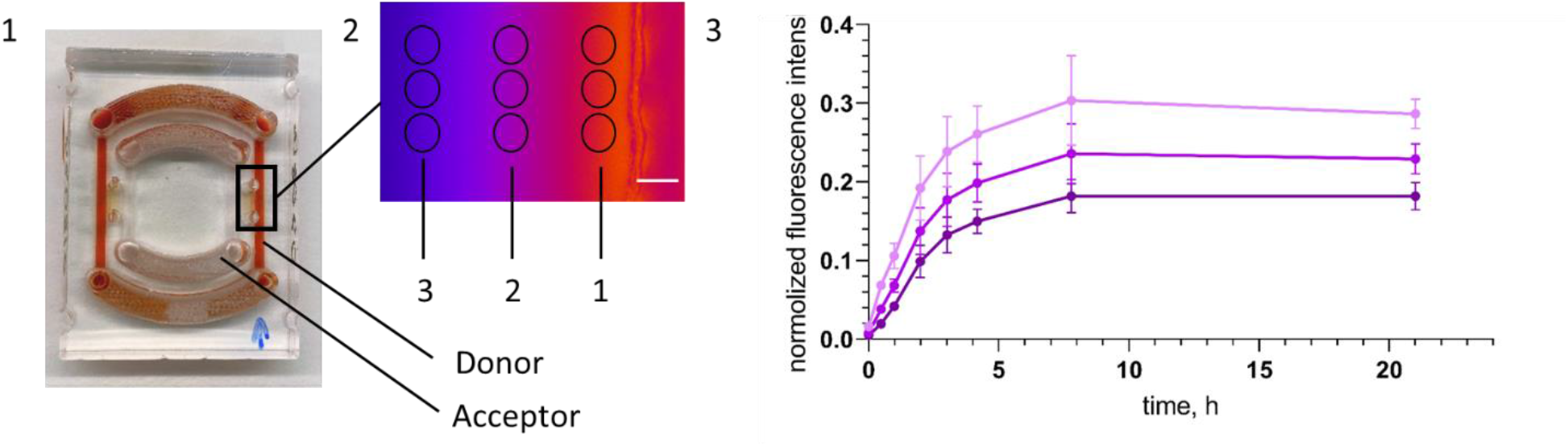
gradient generation analyzed using Dextran-diffusion in the rOoC system: a) rOoC filled with red food dye to indicate Donor side. 2) False-colored image of fluorescence intensity (red: high, blue: low) and three measurement spots (1, 2, 3) indicated (scale bar: 100 um). 3) Relative fluorescence intensity of 70 kDa Dextran at the three spots normed to the donor concentration.

## Appendix B: Tilting function

To derive the formulas for the tilting function, we first recall that the rotation of a vector about the x-axis (roll with angle *γ*) and the y-axis (pitch with angle *β*) is given by the rotation matrix

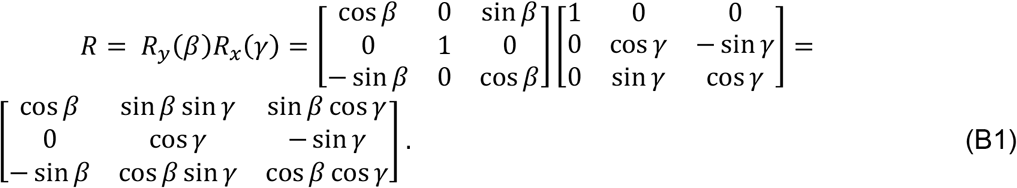

If the upward unit normal vector on the platform base, *n*, rotates with a constant tilt angle *τ* around *e*_*z*_ = [0,0,1]^*T*^, we have

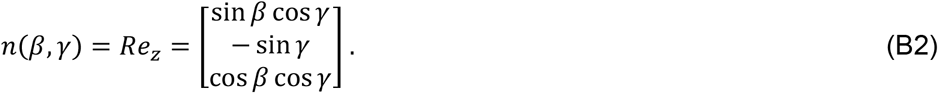

Moreover, we know that the normal vector can be parametrized in term of ϑ(*t*) = 2*πft* with the rotation frequency *f* in Hz and time t in seconds as

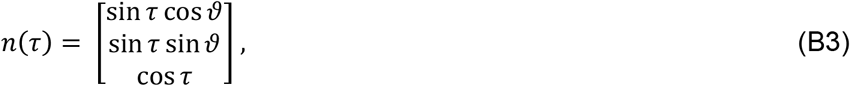

based on the circle parameterization of the rotation:

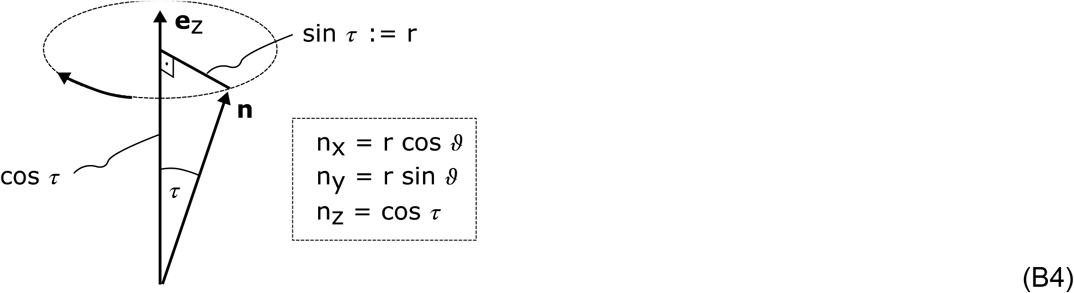

By comparison, we get

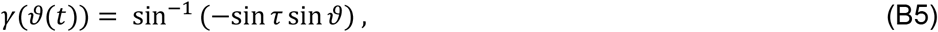

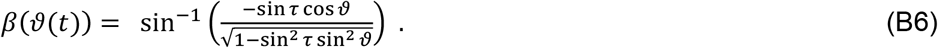

With the remaining relation cos *τ* = cos *β* cos *γ* = *n* ∙ *e*_*z*_, we can easily verify that the tilt angle *τ* is indeed independent of *t*. The pitch and roll angles can also be approximated as

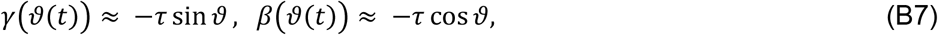

with an approximation error of less than 1% for tilt angles smaller than 20°.

## Appendix C: Model development and technical characterisation

In a rough approximation, the reservoirs can be simplified as rectangular (width b_R_, length l_R_). The height difference within the reservoirs (Δh_R_) and between the in- and outlets of the PCs (Δh_c_) can then be calculated from the pitch and roll β, γ and lengths.

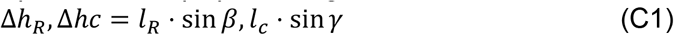

The water table in the reservoir is always in a hydrostatic equilibrium meaning that the liquid level h_i,j_ can be calculated by dividing the volume by the wetted area (A=b_R_*l_R_). In case of the reservoir tilt Δh_R_ is greater than the respective liquid level h_i,j_, the in-/outlet runs dry:

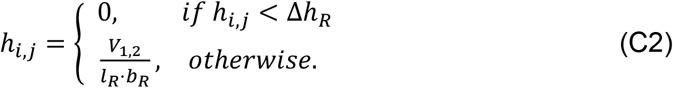

Given these conditions, a simplified model can be defined to estimate the flow rate in the system. The Python code is given in the Suppl. Code 6. It is obvious, that the flow directionality of the system is provided by the shape of the reservoirs and the fact that the tilt Δh_R_ is causing the in-/outlets to run dry. Therefore the volume initially filled in the reservoirs (V_0_) is crucial for the functionality of the device. In the following we are using the following parameters and driving conditions for the modelling and fluidic characterization:

The dynamic viscosity μ for the used media (EGM™-2 BulletKit™, Lonza, Switzerland) is not detailed reported but can be estimated from similar media like DMEM with FBS supplement ^44^. Given the FBS content of 2% for EGM-2, we approximate the viscosity to be1 mPa*s at 20°C and 0.8 mPa*s at 37°C. Table C denotes a different PC cross-section than given in Fig. 1d. The reason behind this is that the optical setup used to measure the flow velocity has a very high focal length thus detecting almost all particles in the PC. The 3d nature of the flow field will lead to a very high tracking error as particle patterns are overlapping each other. To overcome this problem, the PC height for the devices used for the flow studies has been reduced to 0.5 mm. To achieve a similar channel transmissibility σ (approximation error: 1.2%), the PC width was increased to 3.2 mm. In terms of flow rate, this geometry is equal to the cross-section used for the cell experiments but can be measured with the current optical setup without using light-sheet illumination or objectives with a high numerical aperture.

**Table C:** Parameters used for the simplified model

| expression | $l_c$ | $b_c$ | $w_c$ | $l_R$ | $b_R$ | $\alpha$ | $f$ | $\mu$ | $\rho$ |
| --- | --- | --- | --- | --- | --- | --- | --- | --- | --- |
| unit | Mm | mm | mm | mm | mm | ° | rpm | mPa*s | kg/m<sup>3</sup> |
| value | 20 | 3.2 | 0.5 | 20 | 3 | 20 | 3 | 1 | 1001 |

Fig. C1 shows the calculated flow rate and volumes for different initial volumes using the simplified model. Up to a filling volume of 300 μL (excluding the PC volume), the flow is unidirectional as the reservoirs run dry and the whole volume is exchanged. Above this value, a negative flow component is occurring due to liquid remaining on the other side of the tilted reservoir causing a reversed flow at certain positions of the platform. The transported volume is either defined by the amount filled in (V_0_) or the maximum tilt and the reservoir geometry:

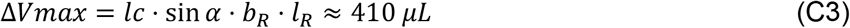

If the reservoirs are filled with a larger volume than ΔV_max_, none of them in-/outlets ever run dry and thus the flow-time curve follows the platform movement (Eq. 1). A higher volume will only add dead volume but not change the flow characteristics as visible in the volume-time-curves for a filling volume of 1.1 mL.

To validate the developed model, we conducted flow measurements with the μPIV setup explained in the Materials and Methods section. The flow was measured in the middle between both reservoirs of the outer PC. A flow-time curve for different initial filling volumes is given in Fig. C2. In contrast to the simplified model, the flow is in reality unidirectional even for higher filling volumes and rises much faster. Moreover, in contrast to Fig. C1, the flow rate is much reduced for a filling volume of 600 μL compared to 500 μL.

Another important aspect is surface effects at the air-liquid interface. When the reservoirs run dry, an air-liquid interface is formed that will present a resistance to the flow according to Eq. 6. To obtain a reasonable value for the critical pressure p_c_, the contact angle has to be measured. We used drops of deionized (DI) water on different substrates of PMMA (Contact Angle Goniometer, Ossila, UK). In Fig. C3/4, the contact angle for a pure PMMA substrate and after exposure to a dose of 1.2 J/cm^2^ UV light is shown. The contact angle is reduced from 80° to 20° and the grade of hydrophilicity depends on the applied UV dose and fades away only slightly over one year (Fig. C5). The critical pressure depends furthermore on the surface tension δ (for cell media values between 50 and 70 mN/m have been reported ^45,46^) and the curvature of the liquid-air interface. As these values are difficult to define exactly, the critical pressure was measured experimentally for different in-/outlet geometries. The experimental setup was as follows:

- A cylindrical reservoir (radius r) is filled with a defined volume V of liquid to obtain a gravity-driven pressure: 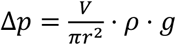.
- The supply reservoir is connected with a PC (width: 1mm, height: 0.5 mm) to a receiving reservoir with different inlet geometries.
- The liquid level is increased until the receiving reservoir is filled. The needed value of Δp is defined as the critical pressure p_c_ for this geometry and cell media.

By using this method, we determined a critical pressure of 40 Pa.

**Fig. C:**
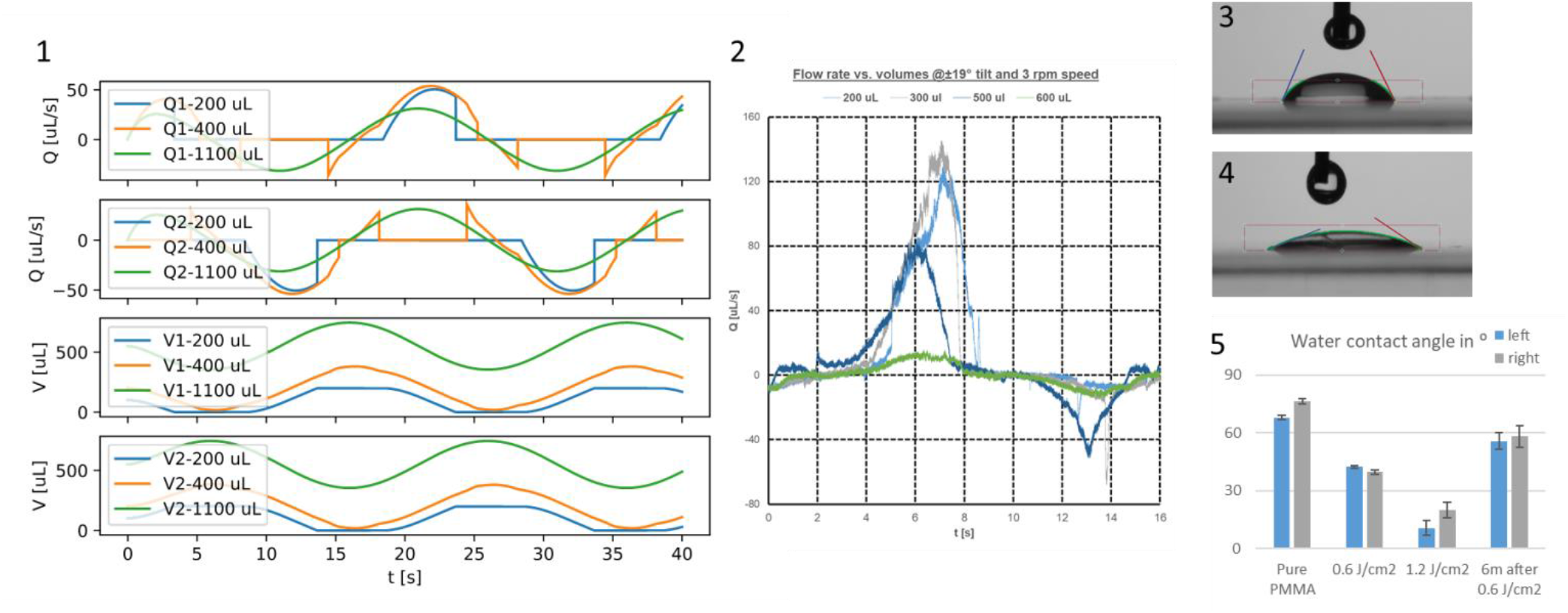
1) Flow rates (Q1/2) and volumes (V1/2) calculated with the simplified model for different filling volumes. 2) Measured flow rates in the outer PC using the μPIV setup for different filling volumes. 3) Contact angle snapshot at pure PMMA. 4) Contact angle snapshot after exposure with 1.2 J/cm^2^ UV light. 5) Contact angle values of DI water on PMMA for two different exposure doses and after 6 months of storage. (n=100, N=3)

## Notes

### Competing Interest Statement

The authors have declared no competing interest.

### Summary of Updates

We include new experimental data for the flow measurement, gradient generation, modeling, and immune experiments, rearranged and reformulated the text and the images to make it easier to understand, removed the APAP-induced liver toxicity and added more information in the supplementary.

