## Supplementary material for "Pump-less, recirculating organ-on-a-chip (rOoC) platform": Suppl. Text

### Channel flow rate, velocity, wall shear stress

#### 1 Fully-developed laminar incompressible flow in a straight channel

Fluid flow of an incompressible ($\rho$ = const.) Newtonian fluid in a channel is described by the Navier-Stokes equations,

$$\rho \frac{\partial \boldsymbol{u}}{\partial t} + \rho \operatorname{div}(\boldsymbol{u} \otimes \boldsymbol{u}) - \operatorname{div}(2\mu \boldsymbol{D}(\boldsymbol{u}) - p\boldsymbol{I}) - \rho \boldsymbol{g} = \boldsymbol{0}, \tag{1a}$$

$$\operatorname{div} \boldsymbol{u} = 0, \tag{1b}$$

with $\boldsymbol{D}(\boldsymbol{u}) = \frac{1}{2}\left(\nabla \boldsymbol{u} + \nabla^T \boldsymbol{u}\right)$ or component-wise in Cartersian coordinates (and simplifying the viscous term using the continuity equation[1]),

$$\rho\left(\partial_t u_x + u_x\, \partial_x u_x + u_y\, \partial_y u_x + u_z\, \partial_z u_x\right) + \partial_x p - \mu\left(\partial_x^2 u_x + \partial_y^2 u_x + \partial_z^2 u_x\right) - \rho g_x = 0, \tag{2a}$$

$$\rho\left(\partial_t u_y + u_x \partial_x u_y + u_y \partial_y u_y + u_z \partial_z u_y\right) + \partial_y p - \mu\left(\partial_x^2 u_y - \partial_y^2 u_y + \partial_z^2 u_y\right) - \rho g_y = 0, \tag{2b}$$

$$\rho\left(\partial_t u_z + u_x \partial_x u_z + u_y \partial_y u_z + u_z \partial_z u_z\right) + \partial_z p - \mu\left(\partial_x^2 u_z - \partial_y^2 u_z + \partial_z^2 u_z\right) - \rho g_z = 0, \tag{2c}$$

$$\partial_x u_x + \partial_y u_y + \partial_z u_z = 0, \tag{2d}$$

with velocity $\boldsymbol{u} = (u_x, u_y, u_z)^T$, pressure $p$, (constant) fluid density $\rho$, (constant) dynamic fluid viscosity $\mu$, gravitational acceleration $\boldsymbol{g} = (g_x, g_y, g_z)^T$, and $\partial_k(\cdot)$ denotes the partial derivative $\partial(\cdot)/\partial k$. Stationary, fully-developed laminar flow corresponds to the assumptions:

$$\partial_t \boldsymbol{u} = \boldsymbol{0}, \tag{3}$$

$$v_x = v_y = 0, \tag{4}$$

where the coordinate $z$ corresponds to the channel axis. That means, we have irrotational, layered, unidirectional flow in the channel direction. From the continuity equation Eq. (2d), we obtain that $\partial_z u_z = 0$. Therefore, Equation (2) can be simplified to a simple Poisson equation,

$$\frac{\partial^2 u_z^2}{\partial x^2} + \frac{\partial^2 u_z}{\partial x^2} = \frac{1}{\mu}\left(\frac{\partial p}{\partial z} - \rho g_z\right) := -\frac{G}{\mu}, \tag{5}$$

valid for *arbitrary* channel cross-sections. To be solvable, we need to provide boundary conditions on the channel wall, which usually is the no-slip boundary condition $\boldsymbol{u} = \boldsymbol{0}$, also referred to as homogeneous Dirichlet boundary condition. $G$ is a constant and measures the driving force.

Wall shear stress vectors are given by the tangential projection of the shear stress tensor onto the channel boundary,

$$\boldsymbol{\tau}_w = \boldsymbol{\sigma n} - (\boldsymbol{\sigma n} \cdot \boldsymbol{n})\boldsymbol{n}, \quad \text{where} \quad \boldsymbol{\sigma} = 2\mu \boldsymbol{D}(\boldsymbol{u}) - p\boldsymbol{I}, \tag{6}$$

where $\boldsymbol{n}$ denotes the outward-point unit normal vector on the channel wall. Wall shear stress (WSS) is then defined as the magnitude of the wall shear stress vector, WSS $:= \|\boldsymbol{\tau}_w\|$.

#### 2 Rectangular cross-sections

In a rectangular cross-section with width $W := 2w$ and height $H := 2h$, $W \geq H$, an analytic solution to Eq. (5) is found by separation of variables, and given in [4] or [3], for $x \in [-w, w]$ and $y \in [-h, h]$,

$$u_z(x, y) = G\frac{h^2}{2\mu}\left[1 - \hat{y}^2 + 4\sum_{k=1}^{\infty} \frac{(-1)^k}{\beta_k^3} \frac{\cosh b_k \frac{x}{h}}{\cosh b_k \gamma} \cos b_k \hat{y}\right] \tag{7}$$

$$\beta_k = (2k-1)\frac{\pi}{2}, \quad k = 1, 2, \cdots,$$

[1] $\operatorname{div}(2\mu \boldsymbol{D}(\boldsymbol{u})) = \operatorname{div}(\mu \nabla \boldsymbol{u})$, due to $\operatorname{div} \boldsymbol{u} = 0$, i.e. for incompressible fluids.

with $\hat{y} = y/h$ (dimensionless height) and $\gamma = w/h$ (aspect ratio). The flow rate is given by [3]

$$Q = G\frac{H^3W}{12\mu}\left[1 - \frac{6}{\gamma}\sum_{k=1}^{\infty}\frac{\tanh\beta_k\gamma}{\beta_k^5}\right] := GLt, \tag{8}$$

where $L$ denotes channel length, and $t := \frac{Q}{GL}$ defines the channel transmissibility. The transmissibility is given in the following for convenience. We also give an approximation for high aspect ratios $W \gg H$ ($\tanh\beta_k\gamma \approx 1$),

$$t = \frac{H^3W}{12\mu L}\left[1 - \frac{6}{\gamma}\sum_{k=1}^{\infty}\frac{\tanh\beta_k\gamma}{\beta_k^5}\right] \approx \frac{H^3W}{12\mu L}\left[1 - \frac{0.63}{\gamma}\right]. \tag{9}$$

The wall shear stress in an axis-aligned rectangular channel allows for a simple expression due to the geometry. The normal vector on the side wall is given by $\boldsymbol{n}_x = (1,0,0)^T$. Since $u_x = u_y = 0$, and with Eq. (7) there is an analytic expression for $u_z$, we have

$$\mathrm{WSS}_x\big|_{x=w} := \mu\frac{\partial u_z}{\partial x} = GH\left[\sum_{k=1}^{\infty}\frac{(-1)^k}{\beta_k^2}\tanh\beta_k\gamma\cos\beta_k\hat{y}\right]. \tag{10}$$

The maximum wall shear stress occurs on mid-height of the channel ($y = \hat{y} = 0$),

$$\mathrm{WSS}_x^{\max} := \mathrm{WSS}_x\big|_{x=w,y=0} = GH\left[\sum_{k=1}^{\infty}\frac{(-1)^k}{\beta_k^2}\tanh\beta_k\gamma\right]. \tag{11}$$

We remark that the terms in brackets in Eqs. (7) to (9) and (11) are dimensionless. Therefore, the quantities such as flow rate and maximum side wall shear stress can be obtained for different channel dimensions by a simple scaling.

### 3 Characteristic numbers

We define the hydrodynamic diameter as $D_{\mathrm{hy}} = 2\frac{HW}{H+W}$. The Reynolds number is given by

$$\mathrm{Re} = \frac{VD_{\mathrm{hy}}\rho}{\mu}, \tag{12}$$

where $V$ is a characteristic velocity. It relates inertial to viscous forces. Pipe flow is assumed to be laminar for $\mathrm{Re} \lessapprox 2300$. We estimate the effect of transient inertial forces in comparison to the viscous forces by computing the Womersley number. Then the Womersley number is given by

$$\mathrm{Wo} = \frac{D_{\mathrm{hy}}}{2}\left(\frac{\omega\rho}{\mu}\right)^{\frac{1}{2}}. \tag{13}$$

For $\mathrm{Wo} < 1$ transient effects are expected to *not* play a significant role for the shape of the velocity profiles (this concerns mostly inertia effect close to the boundary of the channel).

At the entrance of the channel, the flow is usually not fully developed yet. The hydrodynamic entry length $L_e$ necessary to reach fully-developed flow (meaning a profile that deviates less than 1 % from theoretical fully-developed flow) has been estimated in rectangular ducts, e.g. [2], in the form

$$L_e = \Phi(\gamma^{-1})D_{\mathrm{hy}}\mathrm{Re} \tag{14}$$

where $\Phi$ depends on the (inverse) aspect ratio $\gamma^{-1} = H/W$ of the channel. It it between $\Phi(0) \approx 0.01$ (parallel plate solution) and $\Phi(1) \approx 0.075$, with $\Phi(2/3) \approx 0.07$, $\Phi(1/6) \approx 0.025$, see [2, Table 2].

### 4 Channels in this work

The channel dimensions and characteristics numbers of given in Table 1. We estimate the largest flow velocities to be about $0.2\,\mathrm{m\,s^{-1}}$. The frequency $\omega$ is given by the rotation frequency, and an upper bound in practical

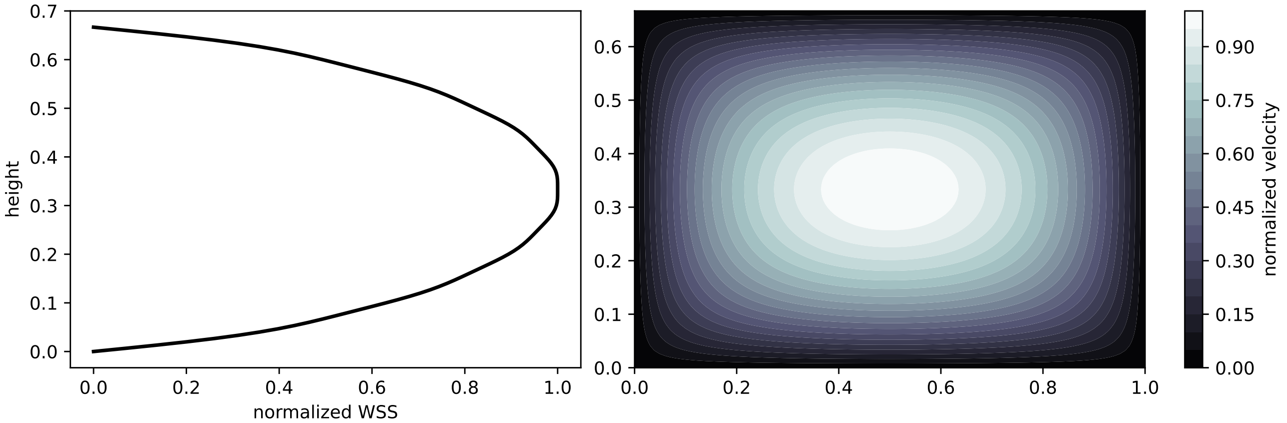


**Figure 1:** WSS distribution over height on the side wall and velocity contours for aspect ratio of Channel 2. WSS, velocity, and width are normalized. Velocity and WSS are computed with Eq. (7) and Eq. (11).

applications is $\omega = 0.17\,\mathrm{s}^{-1}$ (10 rpm). The flow is therefore expected to be laminar in these channels (one of the assumptions for the derivation of the channel transmissibility). Transient effects are expected to *not* play a significant role for the velocity profiles (this concerns mostly inertia effect close to the boundary of the channel). The entry length is relatively large so that the flow is only fully developed in the middle of the channel. This means we make some error in the transmissibility estimate at large velocities, however, this is expected to be less significant than other effects, e.g. capillary effects at the channel entrance. Since, we are interested in wall shear stress (WSS) in the middle of the domain where most of the cells are growing, we expect WSS estimates in the middle of the channel to be sufficiently accurate for the presented analysis.

**Table 1:** Channel dimensions and characteristic numbers for the rectangular cross-section channels analysed with the mathematical model in this work. Reynolds number, Re, Womersley number Wo, and hydrodynamic entry length, $L_e$, are estimates based on the maximal velocities observed in the simulations ($V_{C1s} = 0.1\,\mathrm{m\,s^{-1}}$, $V_{C1l} = 0.12\,\mathrm{m\,s^{-1}}$, $V_{C2s} = 0.1\,\mathrm{m\,s^{-1}}$, $V_{C2l} = 0.12\,\mathrm{m\,s^{-1}}$) and frequency $\omega = 0.17\,\mathrm{s}^{-1}$ which are at the upper end of expected values, and $\rho = 1 \times 10^3\,\mathrm{kg\,m^{-3}}$, $\mu = 1 \times 10^{-3}\,\mathrm{Pa\,s}$.

| | $H$ | $W$ | $L$ | $D_{\mathrm{hy}}$ | $H \times W \times L$ | $\mathrm{Re}_{\max}$ | $\mathrm{Wo}_{\max}$ | $L_{e,\max}$ |
|---|---|---|---|---|---|---|---|---|
| Channel 1 (short) | 0.5 mm | 3.2 mm | 9.8 mm | 0.86 mm | 15.68 µL | 85 | 0.18 | 1.8 mm |
| Channel 1 (long) | 0.5 mm | 3.2 mm | 16.4 mm | 0.86 mm | 26.24 µL | 100 | 0.18 | 2.2 mm |
| Channel 2 (short) | 0.8 mm | 1.2 mm | 9.8 mm | 0.96 mm | 9.41 µL | 100 | 0.20 | 6.7 mm |
| Channel 2 (long) | 0.8 mm | 1.2 mm | 16.4 mm | 0.96 mm | 15.74 µL | 115 | 0.20 | 7.7 mm |

#### 4.1 Transmissibility

Since the channels have aspect ratios close to 1, the approximate solution does not give good results and we go with an approximation infinite series expansion, and cut the sum after 10 terms. Moreover, transmissibilty depends on the fluid viscosity. For velocity measurements at room temperature and culture fluid, we use $\mu_{20} \approx 1\,\mathrm{mPa\,s}$. For wall shear stress calculations at different operating conditions, the temperature is set at $37\,^\circ\mathrm{C}$, and we use $\mu_{37} \approx 0.8\,\mathrm{mPa\,s}$. Therefore, we report here two transmissibility values: $t_{20}$ and $t_{37}$. Recall that flow rate is given by $Q = GLt$ and the mean velocity can be computed by $\bar{v} = Q/(HW)$. Example: for a pressure drop of 50 Pa (upper estimate of what occurs during operation of the platform) over a channel length $L = 20\,\mathrm{mm}$, that is $G = 2500\,\mathrm{Pa\,m^{-1}}$, and $\mu = \mu_{37}$ we have $\bar{v} = 0.1\,\mathrm{m\,s^{-1}}$ for Channel 2 (long) which is what we assumed as upper limit characteristic velocity in the Reynolds number estimates.

**Table 2:** Channel transmissibilities for $\mu_{20} \approx 1\,\mathrm{mPa\,s}$ and $\mu_{37} \approx 0.8\,\mathrm{mPa\,s}$. The product $tL$ is independent of the channel length and only depends on the cross-sectional dimensions.

| | $t_{20}$ | $t_{37}$ | unit |
|---|---|---|---|
| Channel 1 (short) | $3.0664 \times 10^{-9}$ | $3.8330 \times 10^{-9}$ | $\mathrm{m^3\,Pa^{-1}\,s^{-1}}$ |
| Channel 1 (long) | $1.8323 \times 10^{-9}$ | $2.2905 \times 10^{-9}$ | $\mathrm{m^3\,Pa^{-1}\,s^{-1}}$ |
| Channel 2 (short) | $3.0683 \times 10^{-9}$ | $3.8353 \times 10^{-9}$ | $\mathrm{m^3\,Pa^{-1}\,s^{-1}}$ |
| Channel 2 (long) | $1.8334 \times 10^{-9}$ | $2.2918 \times 10^{-9}$ | $\mathrm{m^3\,Pa^{-1}\,s^{-1}}$ |
| | $t_{20}L$ | $t_{37}L$ | unit |
| Channel 1 | $3.0051 \times 10^{-11}$ | $3.7563 \times 10^{-11}$ | $\mathrm{m^4\,Pa^{-1}\,s^{-1}}$ |
| Channel 2 | $3.0069 \times 10^{-11}$ | $3.7586 \times 10^{-11}$ | $\mathrm{m^4\,Pa^{-1}\,s^{-1}}$ |

##### 4.2 Maximum wall shear stress

The maximum wall shear stress may be more conveniently expressed in terms of the flow rate $Q = GLt \Rightarrow G = Q/(Lt)$, such that it can be computed in a post-processing step from the simulated channel flow rates,

$$\mathrm{WSS}_x^{\max} = \frac{QH}{Lt}\left[\sum_{k=1}^{\infty}\frac{(-1)^k}{\beta_k^2}\tanh\beta_k\gamma\right]. \tag{15}$$

With values in SI units, $\mathrm{WSS}_x^{\max}$ is in units of $Pa$ and for reference, we state that $1\,\mathrm{Pa} \mathrel{\hat{=}} 10\ \mathrm{dyne\,cm^{-2}}$, a unit often used for wall shear stress. Figure 1 can be used to determine the wall shear stress on the channel side on specific height of the channel given $\mathrm{WSS}_x^{\max}$. Continuing the previous Example: for $G = 2500\,\mathrm{Pa\,m^{-1}}$, and $\mu = \mu_{37}$ we have $\bar{v} = 0.1\,\mathrm{m\,s^{-1}}$ for Channel 2, thus $Q = 95\,\mathrm{\mu L\,s^{-1}}$. Using the value of Table 3, we obtain $\mathrm{WSS}_{x,37}^{\max} = 0.74\,\mathrm{Pa} = 7.4\ \mathrm{dyne\,cm^{-2}}$.

**Table 3:** Maximum wall shear stress normalized by the channel flow rate $Q$ for $\mu_{37} \approx 0.8\,\mathrm{mPa\,s}$.

| | $\mathrm{WSS}_{x,20}^{\max}/Q$ | $\mathrm{WSS}_{x,37}^{\max}/Q$ | unit |
|---|---|---|---|
| Channel 1 | $6.17 \times 10^6$ | $4.93 \times 10^6$ | $\mathrm{m^{-3}\,Pa\,s}$ |
| Channel 2 | $9.67 \times 10^6$ | $7.74 \times 10^6$ | $\mathrm{m^{-3}\,Pa\,s}$ |

#### 5 Inertial effects

Sudden increases in the pressure gradient are possible due to the gyroscopic motion of the platform in interaction with the reservoir geometry. Without inertial forces considered in the model, the flow rate can increase infinitely fast. Continuing to assume fully-developed velocity profiles at all times, we can still account for inertial forces needed to accelerate the fluid mass. The assumption on the velocity profile allows us to derived from Eq. (2) a one-dimensional model, in form of a momentum and a mass balance (see e.g.[1]),

$$\rho\frac{\partial Q}{\partial t} + A\frac{\partial p}{\partial z} + \frac{\rho\lambda}{A}Q = 0, \tag{16}$$

$$\rho\frac{\partial Q}{\partial z} = 0, \tag{17}$$

with the cross-sectional area of the channel, $A := HW$, and a friction coefficient $\lambda$. For stationary fully-developed pressure-driven flow ($Q = \text{const.}$), we have

$$-\frac{\partial p}{\partial z} = G = \frac{\rho\lambda}{A^2}Q, \tag{18}$$

and we identify $\rho\lambda A^{-2} \equiv (tL)^{-1}$. Thus we obtain from the momentum balance of the whole channel,

$$\frac{\rho}{A}\frac{\partial Q}{\partial t} - G + \frac{1}{tL}Q = 0. \tag{19}$$

We then continue to discretize in time using an backward Euler scheme for a given pressure gradient $G$,

$$\frac{\rho}{A}\frac{Q^{k+1}-Q^k}{\Delta t} - G + \frac{1}{tL}Q^{k+1} = 0, \tag{20}$$

where $\Delta t$ denotes the discrete time step size. Resolving for the flow rate $Q^{k+1}$ at the current time and defining $\zeta := \rho t L A^{-1}$ (in units of seconds), yields with

$$Q^{k+1} = \frac{\zeta Q^k + \Delta t G L t}{\zeta + \Delta t}, \tag{21}$$

an explicit update for the flow rate in terms the flow rate of the previous time step $Q^k$ and the pressure gradient. Note that we always start simulations with with equilibrium conditions such that $Q^0 \equiv 0$. The equilibrium condition can be obtained by running the simulation with $\zeta = 0$ until $Q^{k+1} - Q^k < \epsilon$, while keeping the platform in the initial position.
