## Supplementary material for "Pump-less, recirculating organ-on-a-chip (rOoC) platform": Suppl. Figures

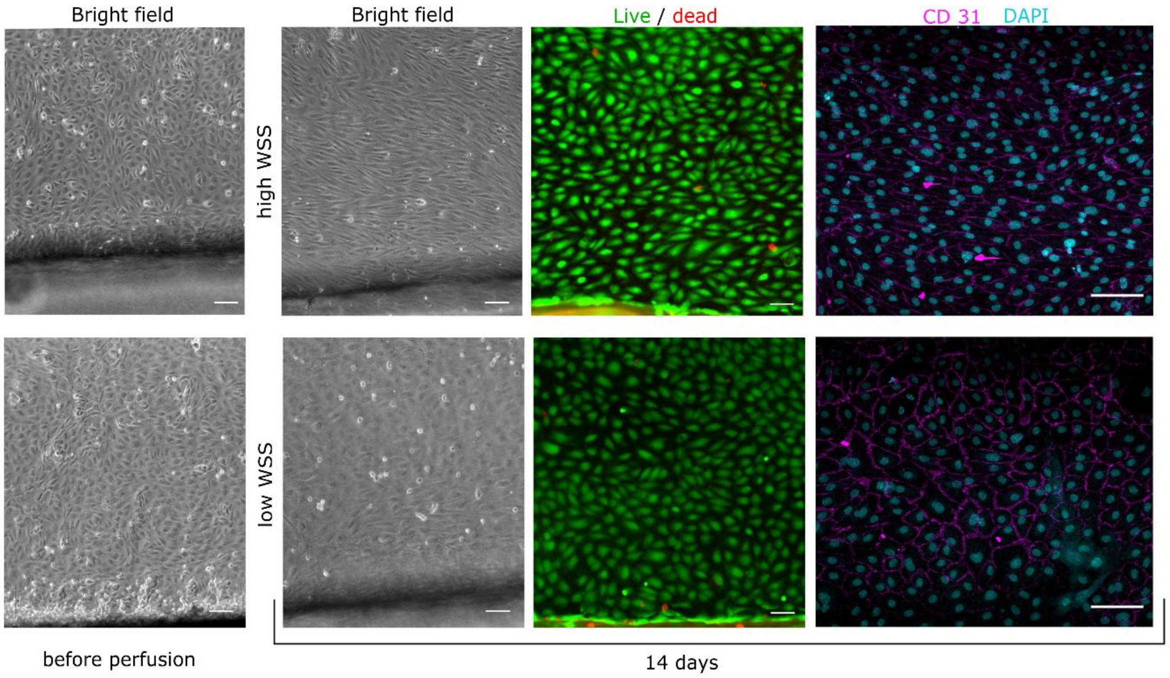


*Suppl. Fig. 1: Long-term culture of HUVEC in the perfusion channels of the rOoC under high and low WSS. Representative Bright-field images, live-dead, and immunofluorescence staining.*

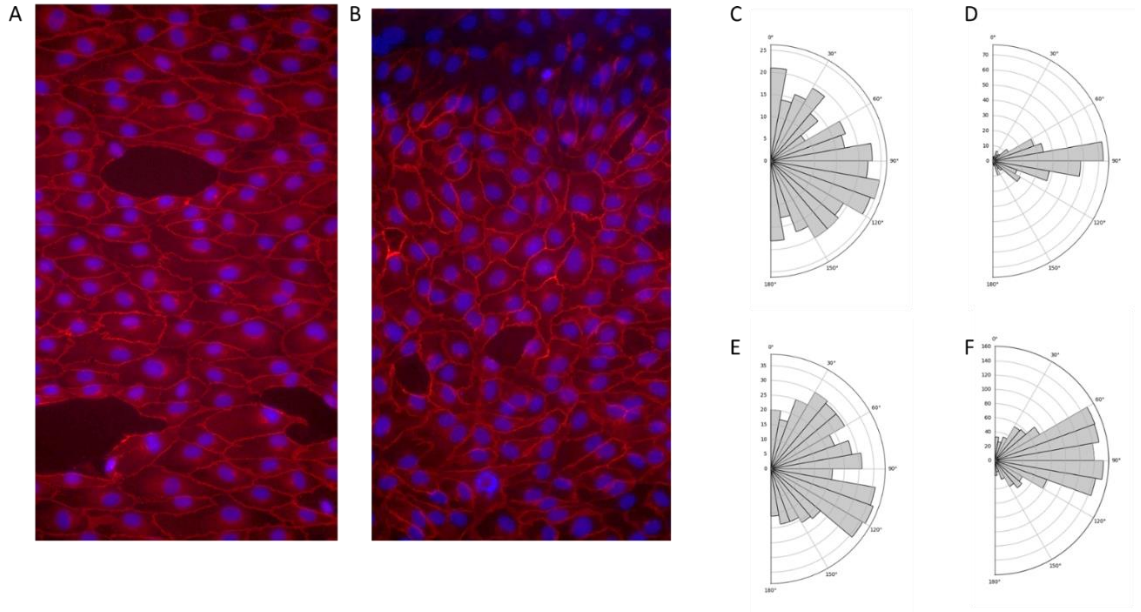


*Suppl. Fig. 2: Alignment of HUVEC cells under different flow regimes: A/B) Fluorescence image of HUVEC cells cultivated in the perfusion channels (red: Actin, blue: Nuclei) in the outer channel (A) and the inner channel (B). C/D: Alignment histogram of nuclei for high flow rate – inner (C) and outer (D) channel. E/F: Alignment histogram of nuclei for low flow rate – inner (E) and outer (F) channel*

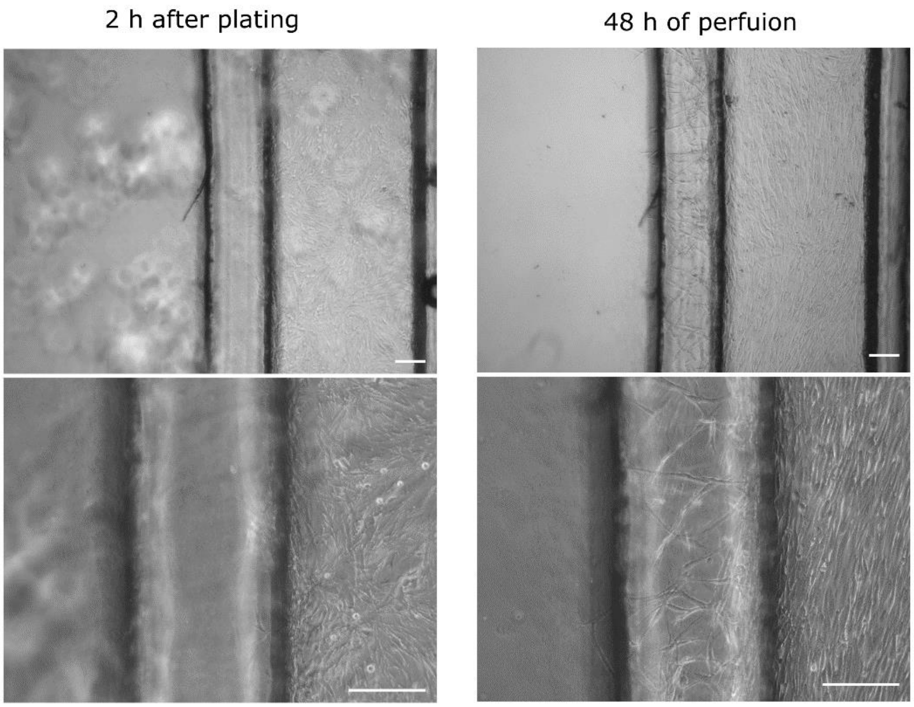


*Suppl. Fig. 3: Human liver endothelial cells (HLEC) in the rOoC. Representative bright field images show alignment in the perfusion channel and sprouting toward the organoid compartment of HLEC after 48 h of perfusion.*

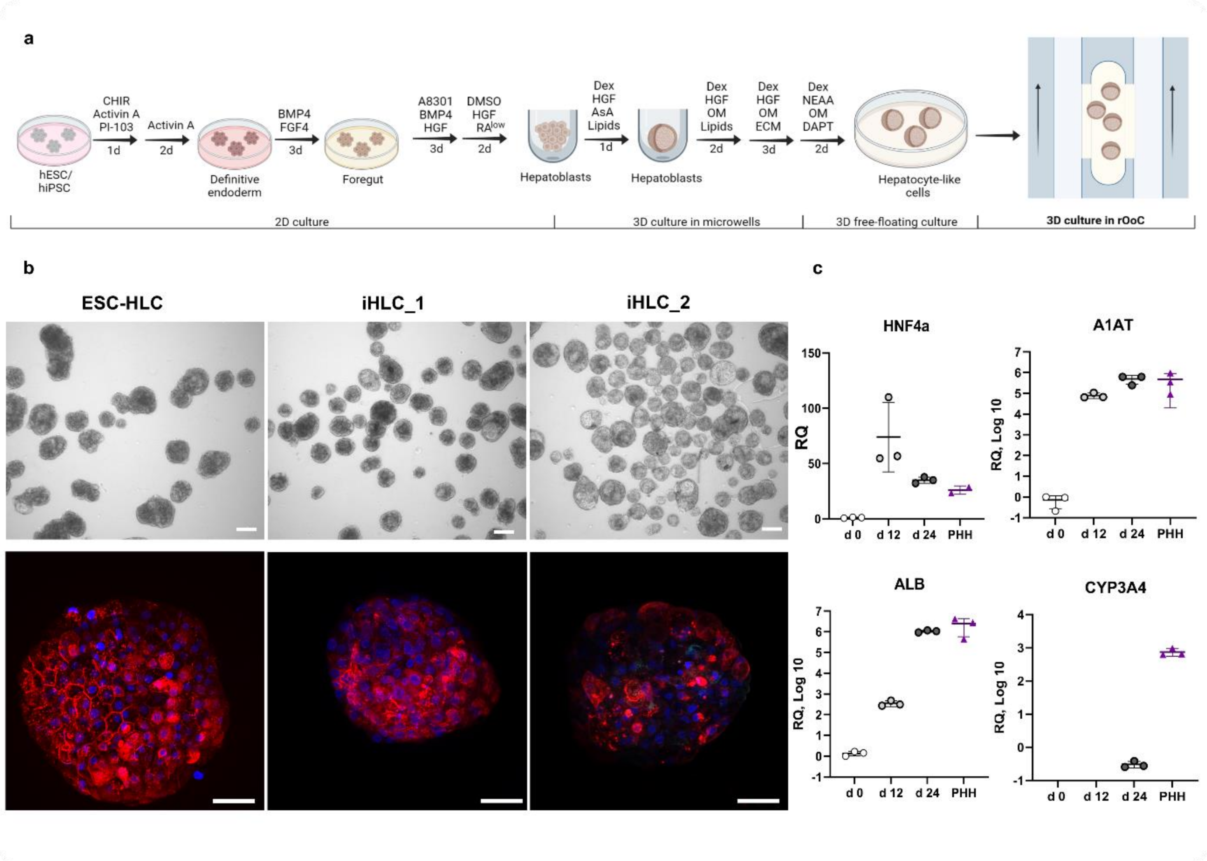


*Suppl. Fig. 4 Differentiation and characterization of embryonic stem cell (ESC) and iPSC-derived human liver cell (HLC) organoids. a) Schematic representation of the used differentiation protocol. b) Representative bright field (upper row) and immunofluorescence (lower row) images of HLC organoids generated from human ESC and iPSC lines. Red – albumin, Blue – nuclei. Scale bar 50 µm. c) Relative expression of selected hepatic markers for three different SC-derived HLC organoids (d 0, d 12, and d 24 of differentiation) compared with primary hepatic organoids (PHH) for n = 3 donors.*

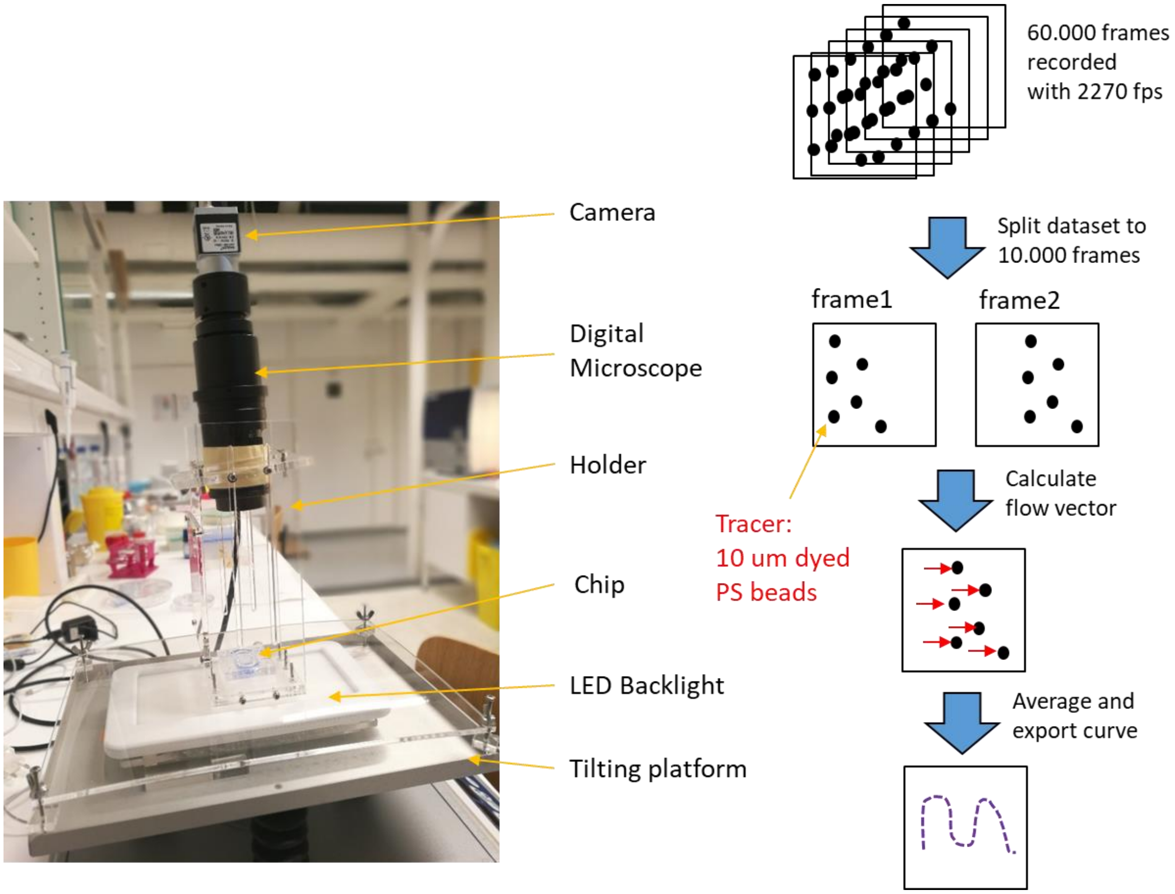


*Suppl. Fig. 5: Optical setup and toolchain for micro-particle-image velocimetry (µPIV) analysis used in this study.*

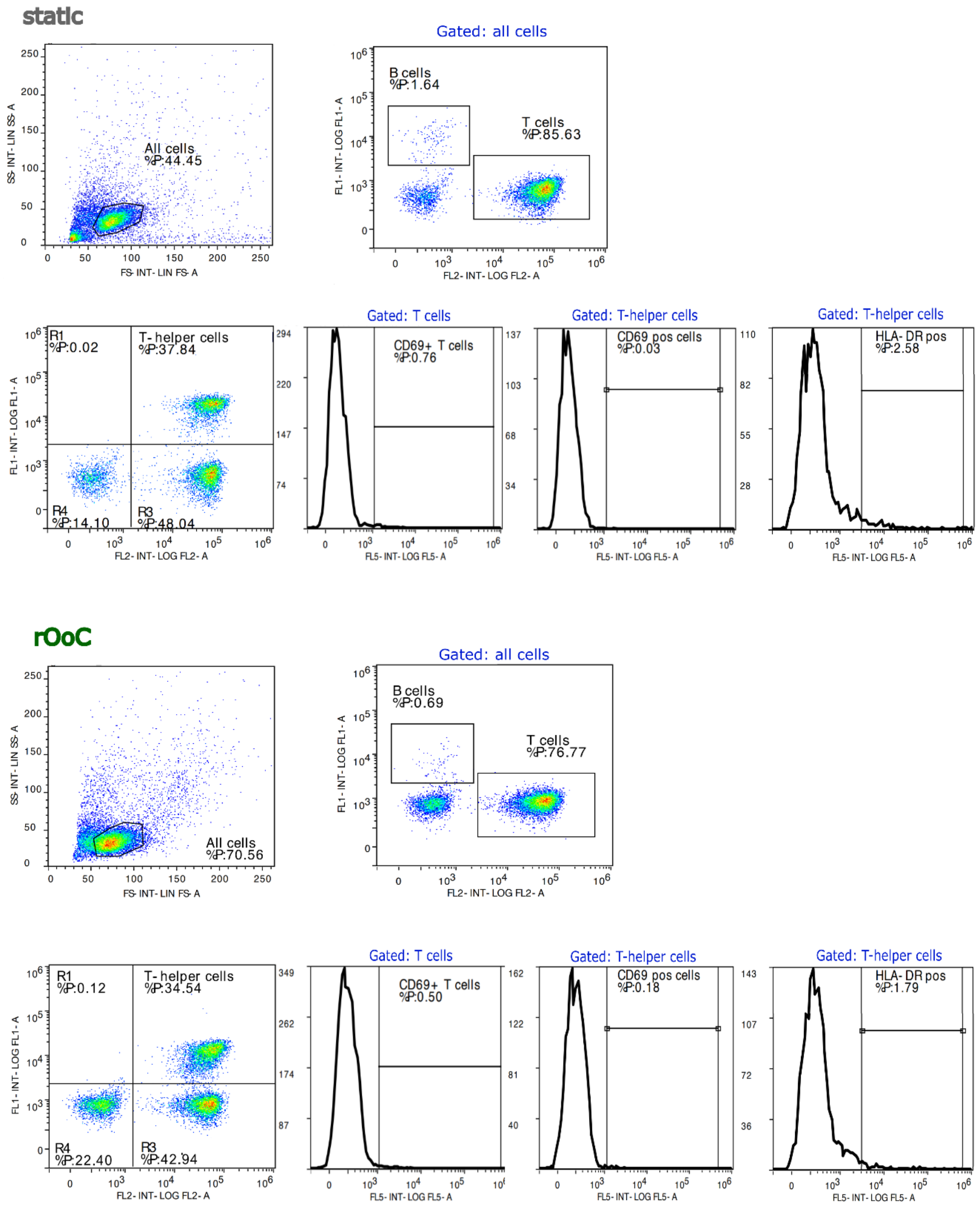


*Suppl. Fig. 6: Gating strategy for human PBMCs. Representative dot blots showing the gating strategy for identifying B cells, T cells, and T-helper cells by flow cytometry. Representative histograms show the percentage of activated T- and T-helper cells under static and fluidic conditions (rOoC) after 24 h. Flow cytometry experiments were replicated for 3 healthy donors.*
