## Supplementary figures and images for "Pump-less, recirculating organ-on-a-chip (rOoC) platform"

### Suppl. Video 2

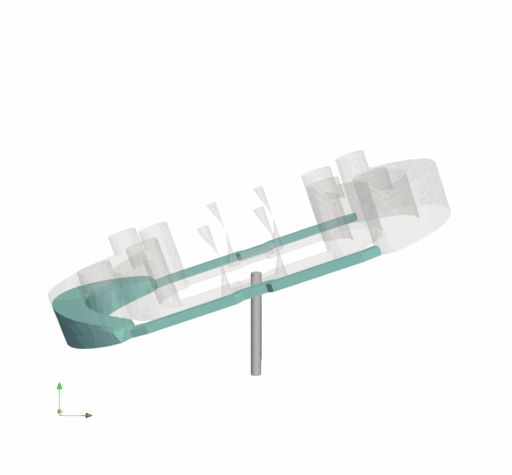
